# Haplotype-phased genome revealed the butylphthalide biosynthesis and hybrid origin of *Ligusticum chuanxiong*

**DOI:** 10.1101/2023.06.13.544868

**Authors:** Bao Nie, Xueqing Chen, Zhuangwei Hou, Cheng Li, Wenkai Sun, Jiaojiao Ji, Lanlan Zang, Song Yang, Pengxiang Fan, Wenhao Zhang, Hang Li, Yuzhu Tan, Wei Li, Li Wang

## Abstract

Butylphthalide, one type of phthalides, is one of the first-line drugs for ischemic stroke therapy, while no enzyme involved in its biosynthesis pathway has been reported. Here, we present the first haplotype-resolved genome of *Ligusticum chuanxiong* Hort., a long-cultivated and phthalide-rich medicinal plant in Apiaceae. Based on comprehensive candidate gene screening, four Fe (II)- and 2-oxoglutarate-dependent dioxygenases (2OGDs) and two CYPs were mined and further biochemically verified as phthalide C-4/C-5 desaturase (P4,5Ds) that converts senkyunolide A to l-*n*-butylphthalide (l-NBP) and ligustilide to butylidenephthalide. The substrate promiscuity and functional redundancy featured for P4,5Ds may contribute to the high phthalide diversity in *L. chuanxiong*. Notably, comparative genomic evidence supported *L. chuanxiong* as a diploid hybrid with *L. sinense* as a potential parent. The two haplotypes demonstrated exceptional structure variance and diverged around 3.42 million years ago (Ma). Our study is an icebreaker for the dissection of phthalide biosynthesis pathway and reveals the hybrid origin of *L. chuanxiong*. These findings will facilitate the future metabolic engineering for l-NBP production and breeding efforts for *L. chuanxiong*.

## Introduction

Dl-3-*n*-butylphthalide (dl-NBP) is one of the first-line drugs for ischemic stroke therapy, which is the commonest type of stroke attacking more than 7.63 million people and causing 3.29 million deaths in 2019^1^. Distinct from the side effects and limitations of recombinant tissue plasminogen activator^2–4^, dl-NBP is prominent for its good neuroprotection and safety in various animal models^5–12^ and was approved for marketing in China in 2004^13^. Moreover, the phase II clinical trial of NBP soft capsules in ischemic stroke patients was started in the United States in 2017^14^. To date, the sales of dl-NBP have reached $1 billion (https://www.cancer.gov/publications/patient-education/takingtime.pdf).

Dl-NBP is a chemically synthetic racemic phthalide (an equal mixture of d-NBP and l-NBP)^15^. Compared with dl-NBP, the naturally distributed l-NBP demonstrated enhanced drug efficiency and reduced side effects. The effects of l-NBP on intracellular cyclic guanosine monophosphate and extracellular nitric oxide levels was antagonized by d-NBP^16^, possibly resulting in a decrease in the drug efficiency of dl-NBP. Moreover, l-NBP is more potent than dl-NBP and d-NBP for antiplatelet^11^, inhibiting apoptosis^17^, hampering TREK-1 (TWIK-related K^+^ channel 1) currents^18^, preventing Alzheimer’s disease^19^, indicating better neuroprotection potential. Thus, pure l-NBP is more promising for drug purposing than dl-NBP^19, 20^. However, l-NBP is less likely to be cost-efficiently produced through chemical synthesis than dl-NBP^21^, and primarily relies on extracting from Apiaceous plants (e.g., *L. chuanxiong*)^22–25^. The low availability hampers the medicinal application of l-NBP. Metabolic engineering of biosynthetic pathways has been proved to be effective and environment friendly in supplying plant secondary metabolites^26, 27^, which may be a promising approach to improve l-NBP production at a relatively low cost.

However, little is known about the phthalide biosynthesis. Previous feeding experiments using ^14^C labeled acetic acid in *Levisticum officinale* revealed that seven C-C units were incorporated into ligustilide (**4** in **Fig. 1a**, an analogue of l-NBP) without rearrangement, which indicated a possible polyketide pathway for the phthalide skeleton forming^28–30^ (**Supplementary Fig. 1**). However, the other possible pathways for the phthalide scaffold formation could not be excluded^31^, such as cyclization of 12-C fatty acid (**Supplementary Fig. 1**). Moreover, it is very challenging to determine the intermediates in polyketide biosynthesis pathway without standard chemicals, though numerous polyketide synthases (PKSs) have been identified^32^. Thus, we shift our attention to the downstream steps. L-NBP belongs to 3-substituted phthalides (type 1 in **Fig. 1b**), which is the most diverse type of phthalides in both plants and fungi^21, 25, 33, 34^. It was most commonly found in the Apiales species **(Fig. 1b**)^21, 25, 34^, particularly in *L. chuanxiong*. Apart from l-NBP, *L. chuanxiong* contains at least 94 other 3-substituted phthalides (**Supplementary Fig. 2**)^35^, among which butylidenephthalide, senkyunolide A and ligustilide are the top three ones (**2–4** in **Fig. 1a**)^35^. They distinguish from l-NBP only in C-4/C-5 or C-3/C-8 olefinic bond (**Fig. 1a**), indicating that they are the likely precursors of l-NBP. Thus, we focus on dissecting the conversion among l-NBP and its three analogues, as it seems a feasible starting point of unraveling the whole biosynthetic pathway.

**Fig. 1.**
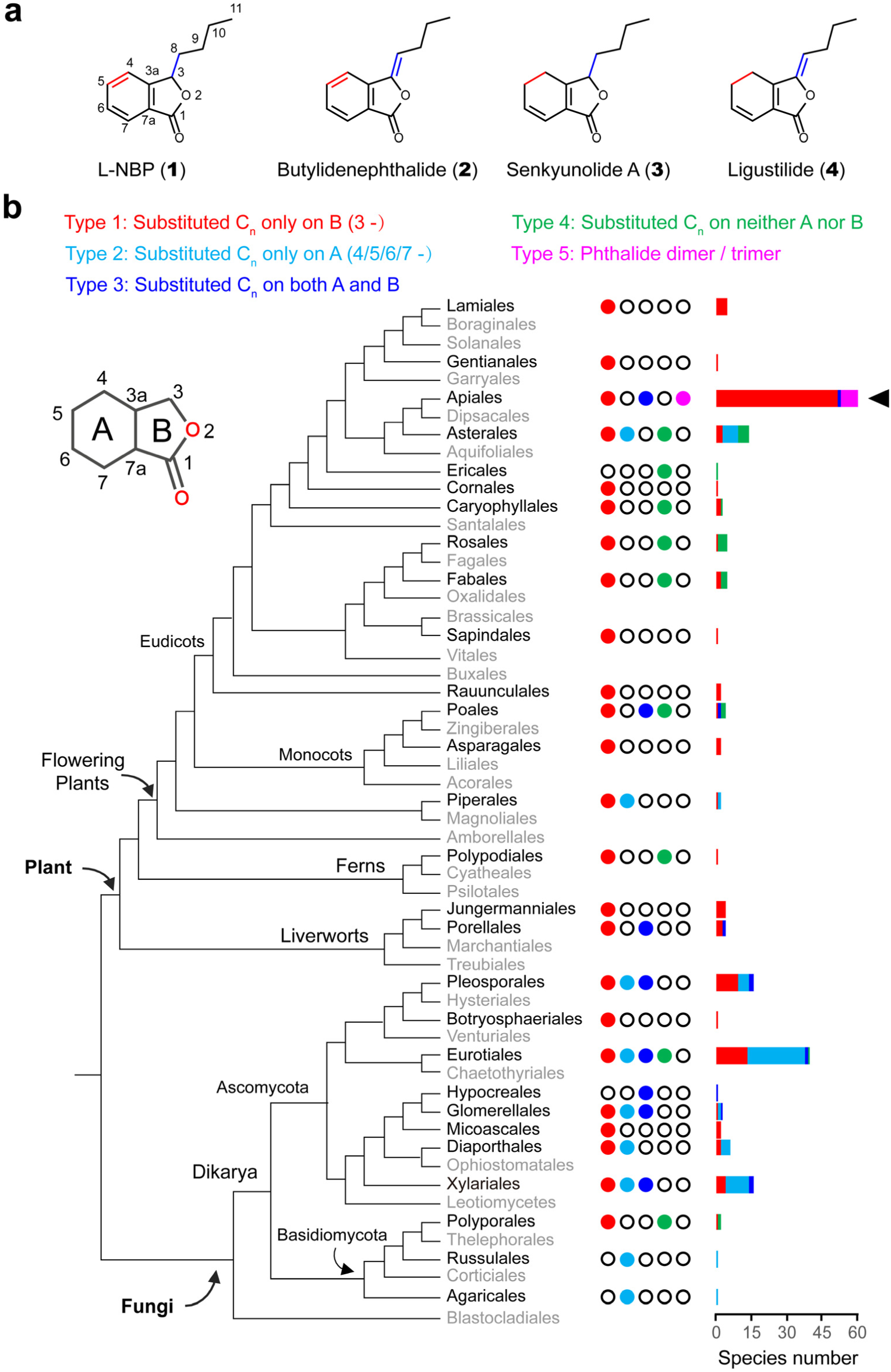
Distribution and diversity of natural phthalides. **a.** The four most abundant mono-phthalides detected in *L. chuanxiong*. The differences among the four phthalides are highlighted with blue (on the side chain) and red (within the ring) colors. **b.** Phylogenetic distribution of natural phthalides, which were divided into five types according to the substituted alkyl position and the polymerization level. Circles filled with colors indicate the presence of the corresponding natural phthalides, and empty circles imply the absence of the corresponding natural phthalides. The horizontal bars denote the number of species in the corresponding order, which have been reported to contain natural phthalides, and the colors of the stacked bars designate the five types of natural phthalides. The information of distribution for phthalides was summarized from several reviews^21, 25, 33, 34^. The four target phthalides in the study all belong to type-1 phthalides (3-substituted phthalides).

*L. chuanxiong,* as an important traditional Chinese medicine (TCM) used to treat heart diseases, has been cultivated for thousands of years through asexual reproduction (vegetatively propagated with expanded stem nodes)^36^ (**Fig. 2a**). To date, no wild population of *L. chuanxiong* has been reported^37, 38^. Thus, *L. chuanxiong* was hypothesized to be a sterile cultivated variety ^36–39^. Despite *L. sinense* being proposed to be a potential wild progenitor of *L. chuanxiong* based on limited evidence^36–39^, this hypothesis need to be further validated based on genomic evidence. The high richness of phthalide and the interesting biological background as a TCM make *L. chuanxiong* as an ideal system to investigate the l-NBP biosynthesis pathway.

**Fig. 2.**
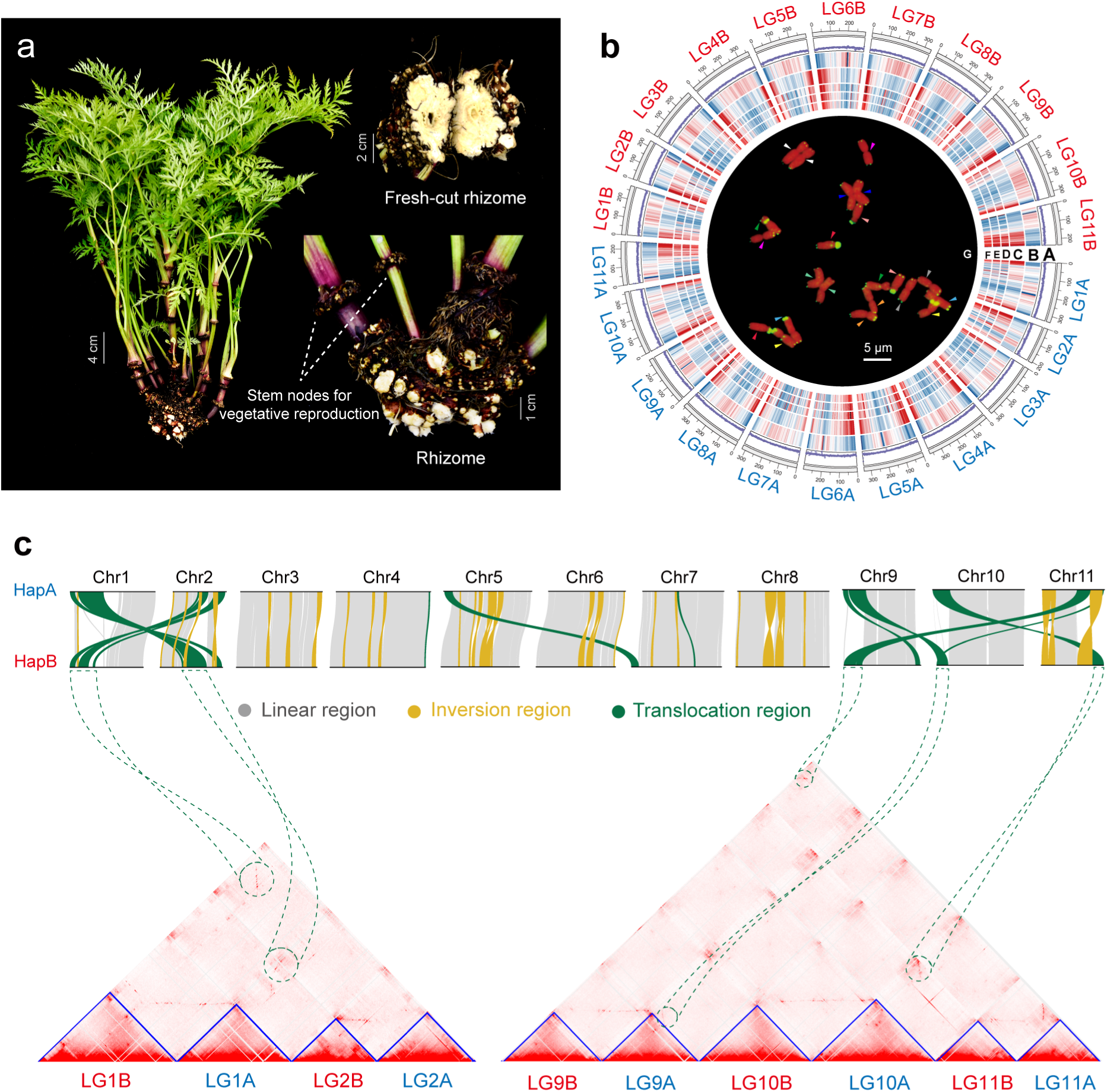
Genomic features of *L. chuanxiong*. **a.** Asexual plant of *L. chuanxiong*. The expanded stem nodes are used for vegetative reproduction. The dried swollen rhizomes are famous TCM “Chuanxiong”. **b.** Genome circos plot and karyotype. (A) GC content, (B) TE density, (C) gene density, (D–F) expression spectrum of different tissues (leaf, stem, rhizome), (G) karyotype. Chromosome number and telomeric repeats distribution (green) of each chromosome were detected by FISH. Arrows in different colors distinguish different homologous chromosome pairs. **c.** SVs larger than 1Mb between two haplotypes. Inversion and translocation are marked with yellow and green, respectively. Hi-C heatmaps show significant interactions (in green circles) between Chr1 and Chr2, as well as among Chr9–11, which provides another piece of evidence for the detected SVs. The resolution of Hi-C heatmap is 2.5 Mb.

Here we present the first chromosome-level and haplotype-phased genome of *L. chuanxiong*. By integrating multi-omics data, biochemical verification, we discovered that four 2OGDs and two CYPs were efficient for catalyzing the conversions among l-NBP and its three analogues. Additionally, we surprisingly found that *L. chuanxiong* was a hybrid based on comprehensive genomic and evolutionary analyses. Limited allele-specific expression (ASE) was detected despite of the high divergence between its two haplotypes. These new findings shed light on the knowledge gap of the phthalide biosynthetic pathway and the origin of *L. chuanxiong*, which paves the way for the metabolic engineering of l-NBP and provides insights into future breeding of *L. chuanxiong* varieties.

## Results

### Genome assembly and annotation of *L. chuanxiong*

Fluorescence in situ hybridization (FISH) revealed 11 pairs of homologous chromosomes in each metaphase root tip cell, which confirmed that *L. chuanxiong* was a diploid species^37, 40^, with *2n* = 2X = 22 (**Fig. 2b**; **Supplementary Fig. 3a**). Further examinations detected asymmetric telomere hybridization signals in at least three homologous chromosome pairs (**Supplementary Fig. 3a**), implying that structural variations (SVs) exist between haplotypes.

Based on 120 Gb PacBio CCS long reads, the genome of *L. chuanxiong* was *de novo* assembled to 7.01 Gb (**Supplementary Table 1–2**), almost twice as the estimate by flow cytometry (3.45 ± 0.38 Gb) (**Supplementary Fig. 3b**), echoing the high genome heterozygosity (4.22%). Given that the phenomenon was previously reported in the highly heterozygous genomes of *Pogostemon cablin* (3.69%)^41^ and *Artermisia argyi* (6.80%)^42^, we inferred that the assembled genome of *L. chuanxiong* contained two divergent haplotypes, which was further validated by Hi-C data. By Hi-C technology, most contigs (96.11% of the genome length) were anchored onto 22 chromosomes, and the predominant “1:1” Hi-C cross-link signals indicated 11 homoeologous chromosome pairs in *L. chuanxiong* (**Supplementary Fig. 4**). Using haplotype-specific k-mers based on SubPhaser^43^, we further phased the genome into two 11-chromosome haplotypes, hereafter referred to as ‘*Lc*HapA’ and ‘*Lc*HapB’ (**Supplementary Fig. 5a–b**). Both haplotype genomes exhibited high completeness from Benchmarking Universal Single-Copy Ortholog (BUSCO) (98.10% for *Lc*HapA; 98.10% for *Lc*HapB) (**Supplementary Table 2**).

Multiple-tissue RNA-Seq data, *ab initio* prediction, and homeolog protein evidence were combined for annotation, totally identifying 101,862 high-confidence protein-coding gene models (52,337 for *Lc*HapA and 49,525 for *Lc*HapB, respectively; **Supplementary Table 2– 3**). Among them, 26,694 pairs were allelic (syntenic and homoeologous) between *Lc*HapA and *Lc*HapB, while the other half was haplotype specific. 10,760,310 repetitive elements were annotated, accounting for 78.45% of the genome. Long terminal repeat (LTR) retrotransposons accounted for the largest proportion (76.02%) of repetitive elements (**Supplementary Table 4**). For both haplotypes of *L. chuanxiong*, expanded gene families mainly enriched in secondary metabolic pathways, while contracted gene families related to immune response, inflorescence meristem maintenance and sexual reproduction (**Supplementary Fig. 6–8**). Consistent with the previous studies^44^, we only detected one whole-genome duplication (WGD) event shared by the Apioideae subfamily, and did not observe recent specific WGD event for *L. chuanxiong* (**Supplementary Fig. 6, 9**).

### Exceptional haplotypic structure variations in *L. chuanxiong*

The high heterozygosity, asymmetric FISH signals, and high proportion of haplotype-specific genes implied great divergence between the two haplotypes of *L. chuanxiong*. Further syntenic analysis revealed substantial SVs between *Lc*HapA and *Lc*HapB (**Fig. 2c**; **Supplementary Fig. 10**). Generally, 39 translocations and 42 inversions larger than 1 Mb were detected, comprising 20.38% of the whole genome, and about half SVs were larger than 5 Mb (**Supplementary Table 5**). On average, each chromosome contained 3.55 inversions and 3.82 translocations larger than 1 Mb. SVs were most remarkable among Chr1, Chr2, Chr 9, Chr 10 and Chr 11, where the largest inversion (∼35 Mb) and translocation (∼37 Mb) occurred (**Fig. 2c**).

Such scale and number of haplotypic SVs were exceptional. We compared the genomic percentage of SVs of *L. chuanxiong* with two other diploid species with either moderate or high heterozygosity (0.69% for *Manihot esculenta* and 1.28% for *Malus pumila*)^45, 46^, two alloploid species (*Perilla frutescens* and *Miscanthus lutarioriparius*)^47, 48^, and a diploid hybrid (*Juglans regia* × *Juglans microcarpa*)^49^ (**Supplementary Fig. 11**). We found that the haplotypic SVs only accounted for 0.12%–0.16% of the genomes in the two diploid species, which was 127–170-fold lower than that in *L. chuanxiong*. In contrast, the inter-subgenome SVs in hybrid or alloploid species, indeed representing interspecific SVs, made up 3.67%– 20.83% of the genome, which were more comparable with the genomic percentage of haplotypic SVs in *L. chuanxiong.* These results jointly suggest the SVs between *Lc*HapA and *Lc*HapB are more prevalent than expected for the intraspecific level.

### Hybrid origin of *L. chuanxiong*

The long history of asexual reproduction of *L. chuanxiong* and the large genetic differentiation between its haplotypes inspired us to speculate that *L. chuanxiong* might be derived from interspecific hybridization instead of solely domesticating from its potential wild progenitor, *L. sinense*^36–39^. To test this hypothesis, we incorporated *L. sinense* into comparative genomic analyses. Given the absence of the genome of *L. sinense*, we *de novo* assembled its transcriptome and obtained 86,244 unigenes (**Supplementary Table 6**).

In order to evaluate the genetic distance among *Lc*HapA, *Lc*HapB and *L. sinense*, we conducted pairwise comparisons among their coding sequences (CDS) from three perspectives: SNPs (single nucleotide polymorphisms), identity, and mapping coverage/quality. First, with *Lc*HapB as the reference, we detected less SNPs for *L. sinense* than *Lc*HapA (**Fig. 3a**). Second, we found that *Lc*HapB and *L. sinense* shared the highest mean identity (99.97%) of orthologous CDS, while *Lc*HapB and *Lc*HapA shared the lowest (99.42%) (**Fig. 3b**). Third, we mapped the RNA sequencing (RNA-Seq) reads of *L. sinense* to each haplotype, and found overall higher mapping rates (88.84% vs. 45.94%) and mapping quality (12.69 vs. 7.18) for *Lc*HapB (**Supplementary Fig. 12**). Taken together, the CDS of *Lc*HapB was more identical with *L. sinense* than *Lc*HapA, indicating a closer genetic relationship between *Lc*HapB and *L. sinense*.

**Fig. 3.**
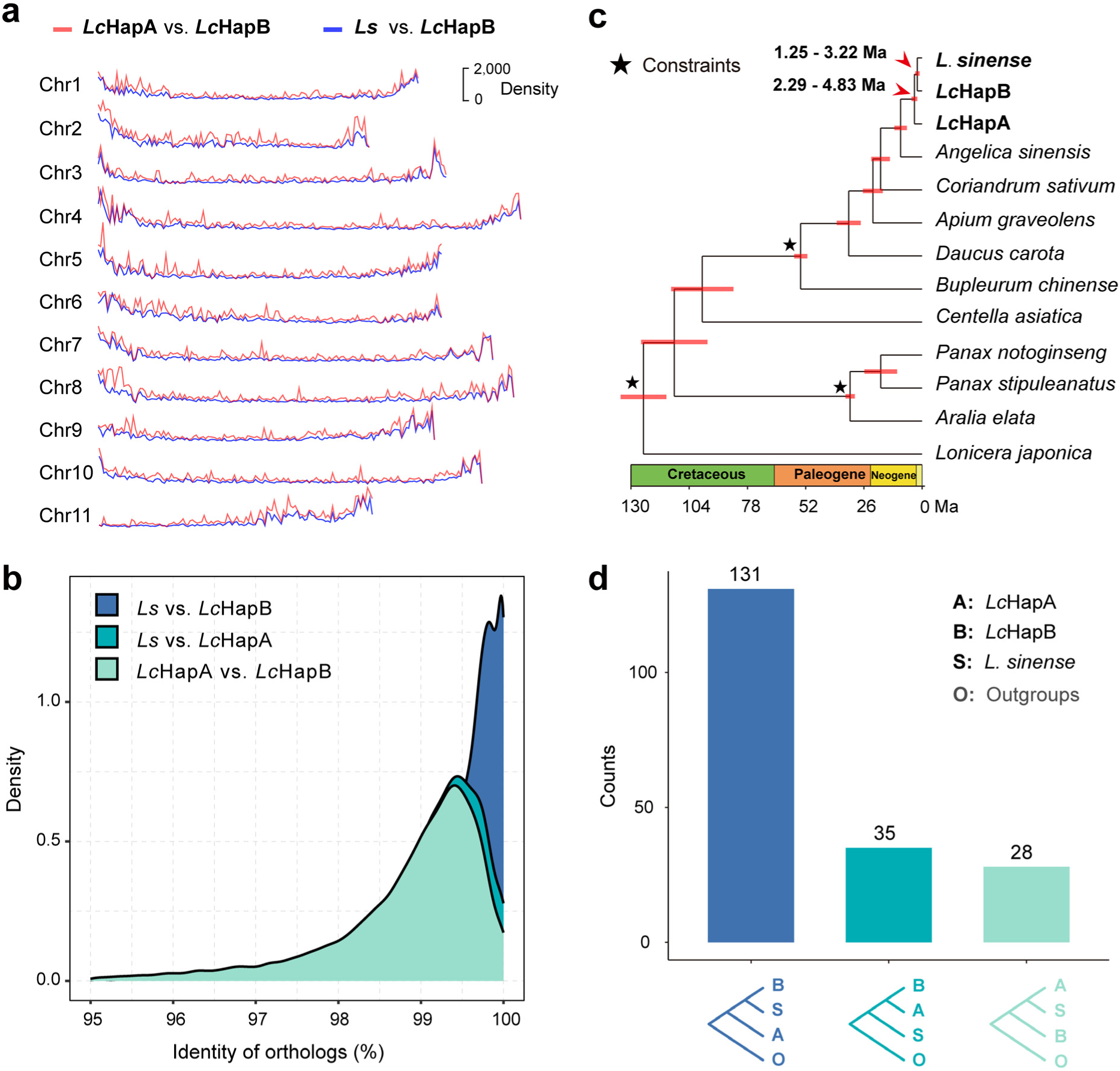
Genetic divergence among haplotypes of *L. chuanxiong* (*Lc*HapA/B) and *L. sinense* (*Ls*). **a.** Comparison of SNP density within 100 kb non-overlapped sliding windows among *Lc*HapA, *Lc*HapB, and *Ls*. **b.** The identity of orthologous CDS among *Lc*HapA, *Lc*HapB, and *Ls*. **c.** The species tree and divergence time among *Lc*HapA, *Lc*HapB, and *Ls*. The constraints for molecular clock calibration are marked with stars. Except that the stem node of *C. sativum* have a bootstrap support of 60, any given node has a support of 100. **d.** Number of gene trees for each identified phylogenetic relationship among *Lc*HapA, *Lc*HapB, and *L. sinense*.

Alternatively, we constructed the phylogenetic trees with 194 single-copy orthologs shared by *Lc*HapA, *Lc*HapB, *L. sinense*, and ten other species from Apiales and Dipsacales (**Supplementary Table 7**). The species tree strongly supported that *Lc*HapB shared the most recent ancestor with *L. sinense* instead of *Lc*HapA with bootstrap support value of 100 (**Fig. 3c**). Specifically, 131 (67.53%) gene trees supported a sister relationship between *Lc*HapB and *L. sinense*, while only 38 (18.04%) presented a closer relationship between *Lc*HapA and *Lc*HapB (**Fig. 3d**; **Supplementary Fig. 13**). Moreover, the molecular clock estimated an earlier divergence age between *Lc*HapA and *Lc*HapB (∼3.42 Ma) than that between *Lc*HapB and *L. sinense* (∼2.12 Ma) (**Fig. 3c**). Thus, the substantial genetic divergence observed between *Lc*HapA and *Lc*HapB might partially represent variation between parental species of the two haplotypes spanning 3.42 million years. In summary, the collective evidence confirmed the diploid hybrid nature of *L. chuanxiong*. *Lc*HapB may be the descendant of *L. sinense* or its relative species, while *Lc*HapA originated from a more distant species that need to be further investigated.

### Balanced allele-specific expression (ASE) in *L. chuanxiong*

Although some level of imbalance between the gene expression of parental alleles in the hybrid has been revealed by several studies^50–52^, our data showed a non-bias pattern of the overall expression between *Lc*HapA and *Lc*HapB (**Fig. 4a**). To examine the allele-specific expression (ASE) of *L. chuanxiong*, we compared the expression level of all 26,694 syntenic and homoeologous gene pairs between *Lc*HapA and *Lc*HapB. We detected ASE in at least one tissue between 4,381 (16.41%) allele pairs, while most alleles (> 70%) did not exhibit significant ASE towards either haplotype in any tissue (**Fig. 4b**). Meanwhile, the ASE genes (ASEGs) towards each haplotype (A-bias or B-bias genes) were roughly even (2,224 vs. 2,157) (**Fig. 4b**; **Supplementary Fig. 14**). These results indicate an overall balanced gene expression between the two haplotypes of *L. chuanxiong*, with limited level of ASE.

**Fig. 4.**
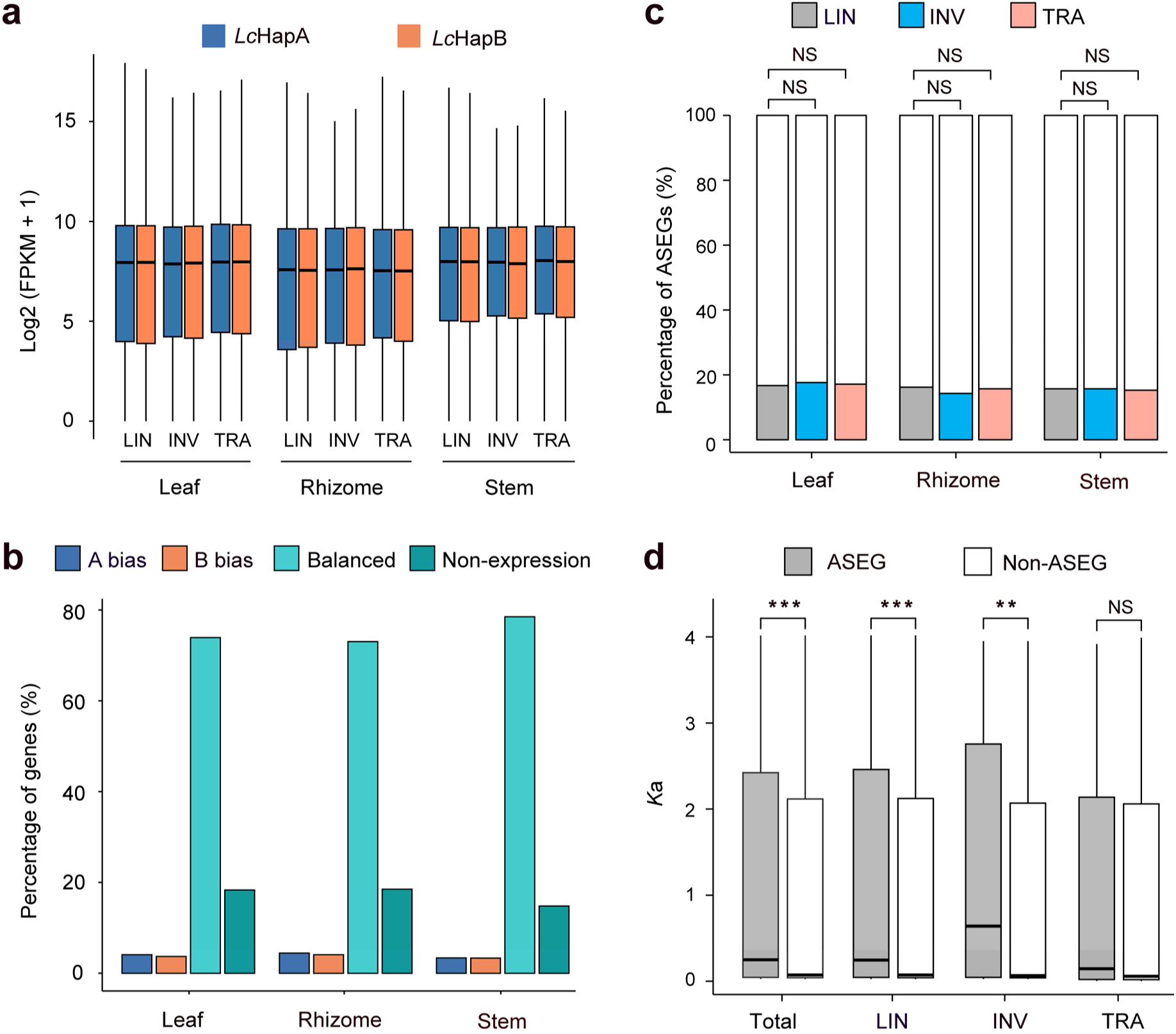
Allele-specific expression (ASE) between the *Lc*HapA and *Lc*HapB of *L. chuanxiong*. **a.** Comparison of gene expression between *Lc*HapA and *Lc*HapB in different SV and tissue types. It is insignificant for all compared groups. **b.** Percentage of genes among different ASE types across three tissues. Alleles were classified into four types according to the expression imbalance between two haplotypes: (1) bias towards *Lc*HapA (A-bias, in blue), bias towards *Lc*HapB (B-bias, in orange), balanced expression between *Lc*HapA and *Lc*HapB (in light green) and non-expression (in dark green). **c.** Comparisons of the percentage of ASEGs among different SV types in three tissues. **d.** Comparisons of *K*a between ASEGs and non-ASEGs in at least one tissue among different SV types and across whole genome. LIN, linear region; INV, inversion; TRA, translocation; Total, whole genome. * *P* < 0.05; ** *P* < 0.01; *** *P* < 0.001; NS, *P* > 0.05.

We further investigated the distribution and functions of ASEGs. Despite exceptional haplotypic SVs in *L. chuanxiong*, we found no significant difference for the proportion of ASEGs between SV and non-SV regions in any tissue (Fisher’s exact test, *P* > 0.05; **Fig. 4c**; **Supplementary Table 8–10**), suggesting that SVs did not affect ASEG distribution. Furthermore, we evaluated the evolutionary rate of ASEGs, and found that ASEGs showed significant higher *K*a (nonsynonymous substitution rate) and *K*s (synonymous substitution rate) than non-ASEGs (Student’s t test, *P* < 0.05; **Fig. 4d**; **Supplementary Fig. 15a–b**). This observation indicates that the evolutionary rate of ASEGs is higher. These fast-evolving ASEGs were mainly enriched in ATP-binding GO term (**Supplementary Fig. 15c–d**) for both A-bias and B-bias genes.

### The inferred scheme of phthalide conversion and *in silico* oxidase gene mining

Based on the high-quality assembled and annotated genome, we started to elucidate the conversion processes among l-NBP (**1**), butylidenephthalide (**2**), senkyunolide A (**3**), and ligustilide (**4**) in *L. chuanxiong*. Given that little is known about the conversion directions and connectivity, we conducted an *in vitro* crude protein activity assay to explore the substrate-product relationships. We initially treated each of the four phthalides with the rhizome crude protein extract (CPE) of *L. chuanxiong*. If any significant signal for the other three phthalides was detected by gas chromatography-mass spectrometry (GC-MS), we would determine the conversion direction as from the substrate phthalide towards the newly appeared one(s). As a result, we found high-confidence signals supporting the conversion from senkyunolide A to l-NBP, and from ligustilide towards butylidenephthalide, less confident signals for other potential conversion processes (**Fig. 5a–b**). Therefore, we mainly focused on these two strongly supported steps involving in production of l-NBP and butylidenephthalide.

**Fig. 5.**
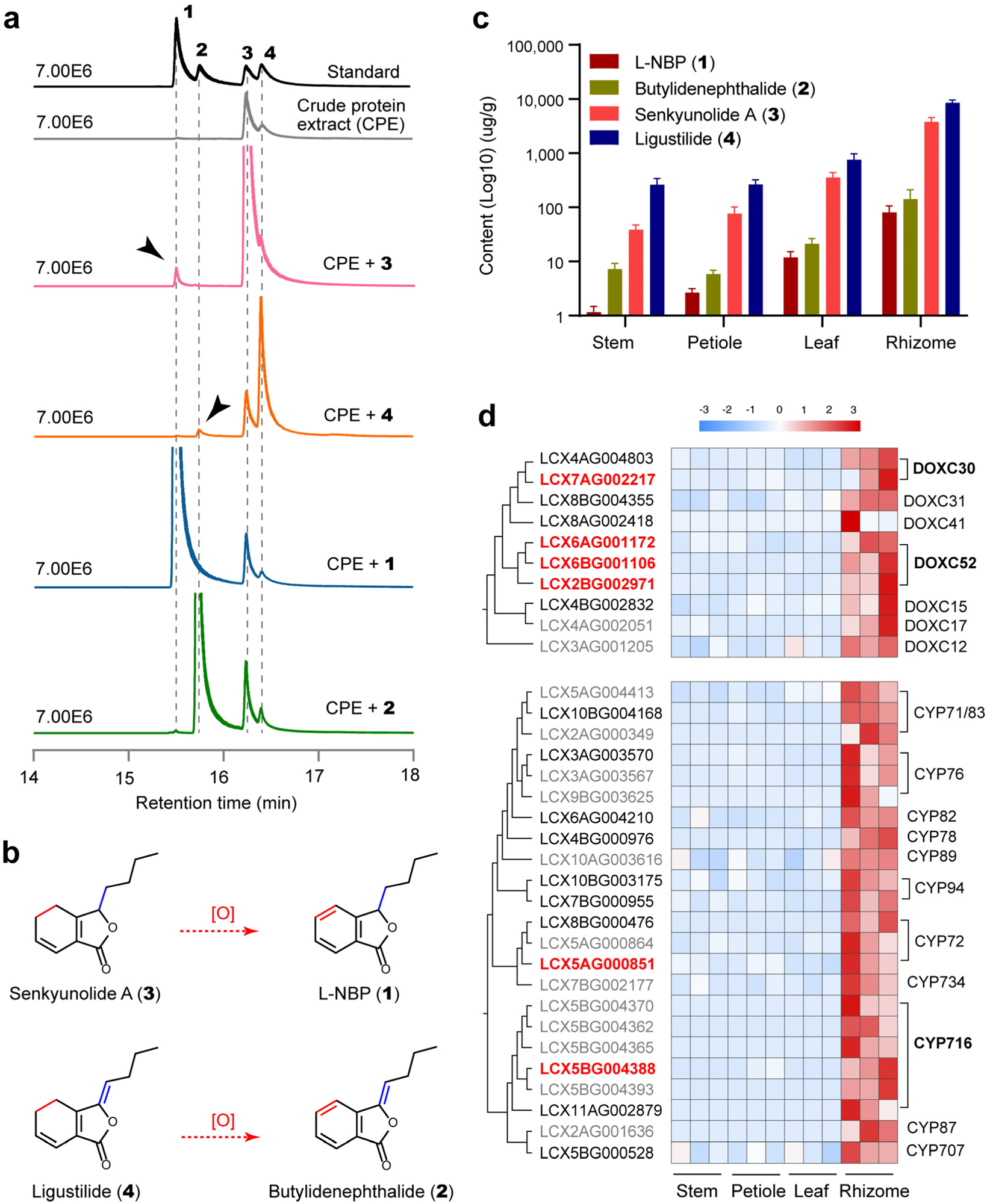
The hypothesis of the conversions among phthalides and mining for the candidate oxidase genes. **a.** GC-MS base peak plot of four phthalides after treating each of them with rhizome crude protein extract (CPE). The alphabets **1–4** represent l-NBP, butylidenephthalide, senkyunolide A, and ligustilide, respectively. **b.** Hypothesis for phthalide conversions inferred from Fig. 5a. The red dashed lines imply confident conversions that were investigated in this study. **c.** Content of phthalide **1–4** from the stem, petiole, leaf, and rhizome extracts of *L. chuanxiong*. Data are presented as the mean values ± standard error of the mean. **d.** Candidate oxidases (2OGDs and CYPs) genes producing l-NBP and butylidenephthalide. The heatmap details the expressional abundance of each candidate gene across three replicates of four tissues. The gradient colour of the scale bar represents relative expression. The gene names on the phylogenetic tree marked with grey indicate genes not successfully cloned; black gene names imply inactive genes; red bold ones show active genes involved in phthalide conversions.

As these two steps are both desaturation processes, we integrated multi-omics data to mine the candidate oxidase genes. Given that *Lc*HapA and *Lc*HapB contained around 50% haplotype-specific genes, we incorporated all genes from both haplotypes into the base gene pool. On ground of the assumption that metabolite accumulation is often positively correlated with the expression of genes involved in metabolic biosynthesis pathway^53^, we initially selected the significantly up-regulated (log2(foldchange) > 1, *P* < 0.05, false discovery rate (FDR) < 0.05) and highly expressed genes (fragments per kilobase of exon model per million mapped fragments (FPKM) > 20) in the rhizome, where the abundance of l-NBP and butylidenephthalide was peaked (**Fig. 5c**). In total, 2,718 genes were chosen (**Supplementary Fig. 16**). Subsequently, we screened out 2,099 and 2,174 genes, whose expression was significantly correlated with the content of l-NBP and butylidenephthalide (Pearson correlation, *P* < 0.05, r > 0.95), respectively. Given that genes of DOXC family in 2OGD superfamily and CYP superfamily typically catalyze oxidation reactions in plant specialized metabolite^54, 55^, we retrieved 14 2OGDs (DOXC family) and 45 CYPs in the 2,099 as well as the 2,174 gene set (**Supplementary Table 11**).

However, it is still experimentally laborious to characterize 59 genes. We conducted phylogenetic analyses to further narrow down the pool of candidate genes. For each DOXC clade or CYP family, we only selected 1–3 highly-expressed genes for cloning (**Supplementary Fig. 17–18**). The final candidate set was limited to nine 2OGDs and 26 CYPs (**Fig. 5d**; **Supplementary Table 11**), among which eight 2OGDs and 11 CYPs were successfully cloned (**Fig. 5d**).

### Four *Lc*2OGDs and Two *Lc*CYPs promoted the phthalide C-4/C-5 desaturations

The enzymatic activities of 2OGDs were verified in *Escherichia coli* system by recombinant protein expression and feeding assays, while the functions of CYPs were characterized in *Nicotiana benthamiana* since we failed to express active recombinant P450 proteins (usually membrane proteins) in *E. coli*. By transiently expressing each cloned 2OGD or CYP, feeding either senkyunolide A or ligustilide as the substrate, and detecting l-NBP and butylidenephthalide as products, we found that four 2OGDs (LCX6BG001106, *Lc*2OGD1; LCX6AG001172, *Lc*2OGD2; LCX2BG002971, *Lc2*OGD3; LCX7AG002217, *Lc*2OGD4), and two CYPs (LCX5AG000851, *Lc*CYP72A1132; LCX5BG004388, *Lc*CYP716E94) could significantly convert senkyunolide A to l-NBP, as well as ligustilide to butylidenephthalide (**Fig. 6a–b**; **Supplementary Fig. 19–20**). Interestingly, we found that fully identical oxidases were involved in the two steps of conversion, which may result from the same desaturation position (C-4/C-5 of phthalide) and similar structure of senkyunolide A and ligustilide. *Lc*2OGD1–2 were homoeologous alleles in Chr6, while *Lc*2OGD3–4 were located in Chr2B and Chr7A. The *Lc*CYPs were distributed in Chr5A and Chr5B. As a result, equal number of phthalide C-4/C-5 desaturases (P4,5Ds) were detected to be effective in *Lc*HapA and *Lc*HapB.

**Fig. 6.**
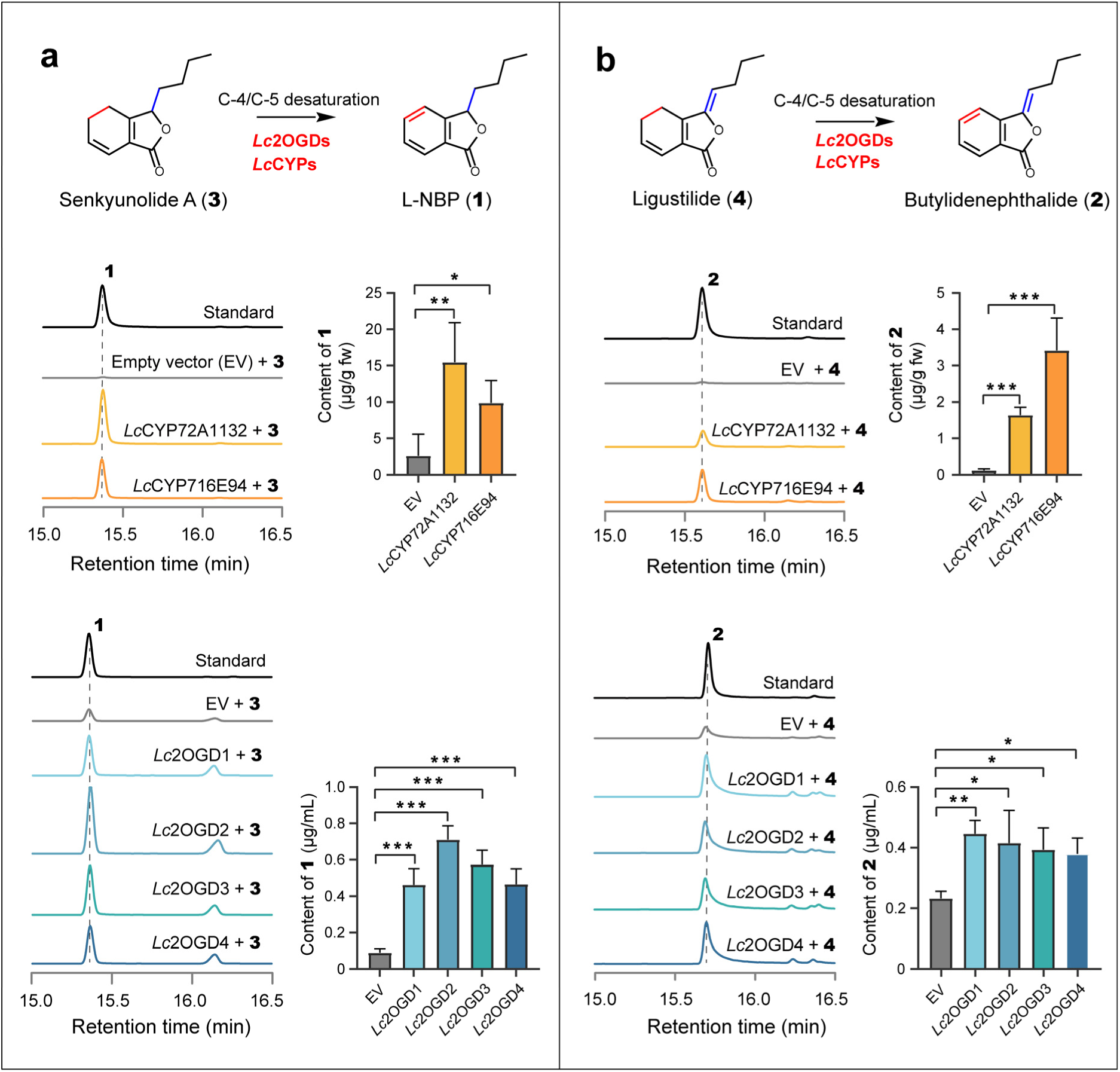
*Lc*2OGD1–4, *Lc*CYP716E94, and *Lc*CYP72A1132 mediate the phthalide conversions. **a.** GC-MS peak plots and the production of l-NBP (**1**) (*m/z* 133) in the indicated combinations of senkyunolide A (**3**) and *Lc*2OGDs or *Lc*CYPs. **b.** GC-MS peak plots and the production of butylidenephthalide (**2**) (*m/z* 159) in the indicated combinations of ligustilide (**4**) and *Lc*2OGDs or *Lc*CYPs. In each panel, the GC-MS peak plot (left) shows the combination of substrate and candidate gene, while the boxplot (right) revealed the content of resulting product in this treatment. All treatments involving different genes and substrates were conducted in triplicate and the error bars represent standard error of the mean. Empty vector (EV) was used as the negative control. All 2OGDs were characterized in *E. coli*, and all CYPs were characterized in *N. benthamiana*. * *P* < 0.05; ** *P* < 0.01; *** *P* < 0.001.

Weak signals of l-NBP or butylidenephthalide were also detected in the negative control groups (empty vector), indicating spontaneous conversion from senkyunolide A to l-NBP, and from ligustilide to butylidenephthalide, whereasprevious studies have demonstrated that these two spontaneous conversions were slow^56, 57^. More importantly, our signals in *Lc*2OGD1–4 and *Lc*CYPs added groups were approx. stronger by 1.63-fold to 27.94-fold than the negative control groups (**Fig. 6a–b**), indicating an efficient and reliable function of these P4,5Ds.

### Redundancy and phylogenetically independent recruitment of the P4,5Ds

As the desaturase function for phthalides has never been reported in 2OGD and CYP superfamily, we further investigated the phylogenetic relationships between P4,5Ds and other 96 characterized 2OGDs and 67 characterized CYPs (**Supplementary Table 12–14**). *Lc*2OGD1–3 belonged to DOXC52 clade, while *Lc*2OGD4 located in DOXC30 (clade names following Kawai et al.^55^) (**Fig. 5d**). DOXC30 comprises of the widely distributed feruloyl-CoA 6’-hydroxylase involved in coumarin biosynthesis pathway^55^ (**Supplementary Fig. 21**; **Supplementary Table 12**). Nevertheless, DOX52 targeted more diverse substrates, including dopamine, 4-hydroxyphenylacetaldehyde^58^, thebaine^59^, and melatonin^60^. In addition, the two *Lc*CYPs belongs to two distant families: CYP72 and CYP716, which are conventionally involved in terpenoid biosynthesis, especially triterpenoid tailoring (**Supplementary Fig. 22– 23**; **Supplementary Table 13–14**). Thus, P4,5Ds are likely to originate from multiple pathways and distinct gene lineages.

Additionally, the functional redundancy was noteworthy for phthalide C-4/C-5 desaturations, involving six genes (*Lc*2OGD1–4, *Lc*CYP72A1132 and *Lc*CYP716E94). This clue prompted us to investigate more potential P4,5Ds by reexamining the candidate CYPs for l-NBP or butylidenephthalide biosynthesis. Interestingly, we found that four more candidate CYP716Es were physically linked to *Lc*CYP716E94. Further examination for adjacent genomic region detected a densely distributed tandem duplication containing 19 copies in *Lc*HapB (∼662 Kb) and nine in *Lc*HapA (∼200Kb). Syntenic analysis for different Apiaceous species showed five CYP716Es in *Bupleurum chinense*, three in *Apium graveolens*, and two in *Coriandrum sativum* within this syntenic region, suggesting that CYP716E subfamily significantly expanded in *L. chuanxiong*, especially in *Lc*HapB (**Supplementary Fig. 24–25**). These copies shared high amino acid sequence identity (57%–96%) with *Lc*CYP716E94, suggesting similar catalytic functions. However, the function of the tandem duplicated copies of *Lc*CYP716E94 warrants experimental verification in the future.

## Discussion

Despite the wide application of NBP for ischemic stroke therapy^13, 20^, no enzyme involved in its biosynthesis has been known to date. The present study is an icebreaker for the dissection of phthalide biosynthesis pathway, where four 2OGDs and two CYPs were proved to be independently recruited for phthalide C-4/C-5 desaturation, converting senkyunolide A to l-NBP and ligustilide to butylidenephthalide in *L. chuanxiong*, respectively. Our data support that the two desaturation processes are mainly attributed to enzymatic rather than spontaneous reactions as previously reported^56, 57, 61^. As senkyunolide A is the second abundant phthalide in *L. chuanxiong* (**Fig. 5c**), ∼47-fold of the content of l-NBP, up-regulation of P4,5Ds *in vivo* would be a promising way to enhance the content of l-NBP in *L. chuanxiong*. Our discovery will facilitate the future metabolic engineering for l-NBP production.

The substrate promiscuity and function redundancy were featured for P4,5Ds. All six P4,5Ds can recognize both senkyunolide A and ligustilide that differs in C-3/C-8 bond, indicating that P4,5Ds are promiscuous to different phthalides and possibly mediate other C-4/C-5 desaturating reactions, such as from senkyunolide F to senkyunolide E (**Fig. 7**), etc. Thus, the substrate promiscuity may greatly contribute to the high phthalide diversity in *L. chuanxiong*, as also reported in other metabolic pathways^54, 62, 63^. Moreover, P4,5Ds are independently recruited from both 2OGD and CYP lineages. To the best of our knowledge, the involvement of both 2OGD and CYP in the same reaction is rarely reported^54, 55, 64, 65^, which may partly owe to our comprehensive screening strategy for desaturases from any potential clade in both 2OGD and CYP superfamily, rather than focus on specific clades as in other studies^66, 67^. Our results unveil the probably underestimated gene function redundancy in plant secondary metabolite biosynthetic pathways and advocate a comprehensive survey of genes from different families with similar functions.

**Fig. 7.**
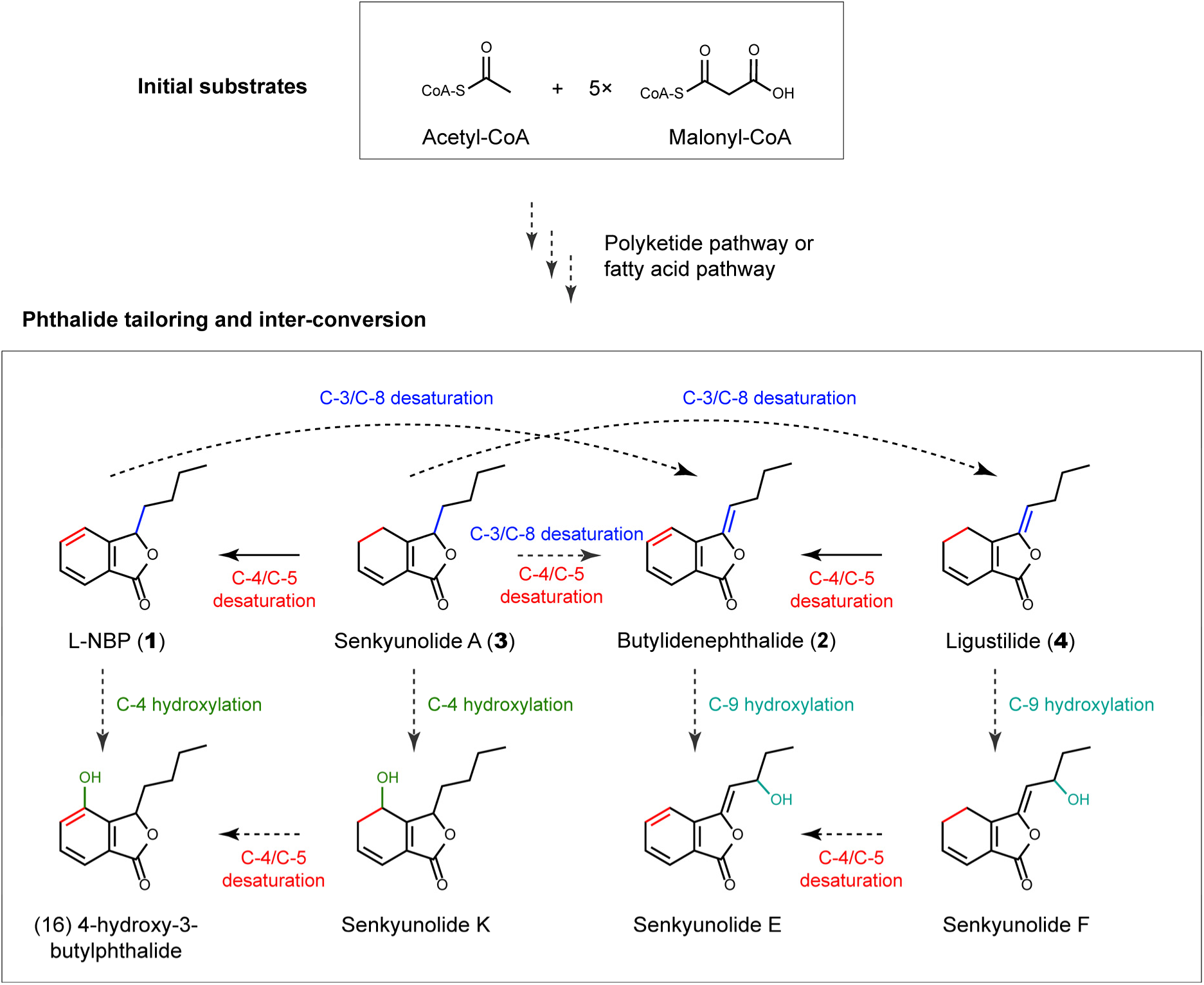
Proposed biosynthetic pathway for l-NBP. The solid lines represent characterized steps in the present study, while the dotted lines represent potential but unresolved steps. Different reaction types were distinguished with different colors. The upstream polyketide or fatty acid pathway was detailed in **Supplementary Fig. 1**.

In addition to the new insights into l-NBP biosynthesis, the firstly reported genome also lifted the curtain of the origin of *L. chuanxiong*. Strikingly, our comparative genomics analyses showed that *L. chuanxiong* was a sterile hybrid from two different diploid progenitors diverged more than 3.42 Ma. One of the parents of *L. chuanxiong* might be or very related to *L. sinense,* which explain their high morphological similarities. However, the other parent is phylogenetically more distant, and awaits further exploration with a denser sampling of related *Ligusticum* species. Due to that *L. chuanxiong* has been vegetatively cultivated for > 1600 year in China^36^, rarely blooming and bearing mature seeds^37, 38^, the taxonomic treatment has been a puzzled problem. Qiu (1979) treated *L. chuanxiong* as a separate species^37^, while Pu (1991) treated it as a cultivated variety of *L. sinense*^38^. Our evidence supports *L. chuanxiong* as a distinct hybrid species, and thus provides a new case for homoploid hybrid speciation (HHS) that is increasingly recognized in recent years^68–70^. Among the HHS examples, the speciation of *Ostrya intermedia* via alternatively inheriting parental isolating allelic genes is impressive^69^. However, *L. chuanxiong* revealed a new isolation mechanism through asexual reproduction that act as an unbridgeable barrier for gene flow between *L. chuanxiong* and its parents. As many diploid hybrid species propagates asexually, such as *Oenothera* (Onagraceae)^71^, *Daphnia pulex*^72^, *Ranunculus*^73^, the loss of sexual reproduction maybe an alternative common HHS mechanism. For *L. chuanxiong*, the sterility may result from several reasons. The substantial chromosomal SVs have been frequently reported to induce higher sterility level because of the inappropriate pairing and separation during meiosis process^71, 74, 75^. Moreover, gene families related to inflorescence meristem maintenance and sexual reproduction were significantly contracted in *Lc*HapA and *Lc*HapB (**Supplementary Fig. 7**), respectively, which may limit the overall ability for flowering of *L. chuanxiong*.

Although the two divergent haplotypes show substantial variation at both sequence and structure level, they generally remain balanced in gene expression and seem to contribute equally to phthalide C-4/C-5 desaturation. The six validated P4,5Ds, along with other candidate genes, were evenly distributed on *Lc*HapA and *Lc*HapB (**Fig. 4d**). Specifically, *Lc*2OGD1 (LCX6BG001106) and *Lc*2OGD2 (LCX6AG001172), one pair of alleles, showed likely equivalent enzymatic efficiency in catalyzing phthalide C-4/C-5 desaturation (**Fig. 6**). Although our discoveries of P4,5Ds represent a small portion of the complete biosynthetic pathway of l-NBP, the possibility that P4,5Ds are not dominant on solo-haplotype but from both haplotypes indicates that the higher content of l-NBP in *L. chuanxiong* than its potential parent *L. sinensis* (**Supplementary Fig. 26**) might benefit from the effect of heterosis on the content of secondary metabolites^76^. Thus, it highlights the significance to further investigate the likely parental species of *L. chuanxiong* with a comprehensive sampling of species in *Ligusticum*, and design artificial interbreeding experiments to evaluate the effect of heterosis on the content of significant secondary metabolites, such as l-NBP, in the future study.

In conclusion, our study provided crucial genomic information of *L. chuanxiong*, firstly identified six P4,5Ds and revealed the hybrid origin of *L. chuanxiong*. These new findings will greatly facilitate the future metabolic engineering for l-NBP production and point out the direction for future breeding of *L. chuanxiong* varieties. However, several questions regarding the phthalide biosynthesis and origin of *L. chuanxiong* have yet to be answered. In addition to phthalide C-4/C-5 desaturations, there exists various oxidatively tailoring processes for phthalide^57, 61^ (**Fig. 7**). More importantly, the dissection of phthalide scaffold forming is crucial for fully resolving the phthalide biosynthetic pathway and for future metabolic engineering (**Supplementary Fig. 1**). Additionally, our study leaves a gap to fill, to investigate the most likely parents of the hybrid, in our future research.

## Methods

### Chemical Reagents

All standard chemical reagents were purchased from commercial vendors unless noted otherwise. The following phthalides were obtained from commercial sources: l-3-*n*-butylphthalide (l-NBP; Chengdu, Chengdu must bio-technology CO., LTD), butylidenephthalide (Chengdu, Chengdu must bio-technology CO., LTD), senkyunolide A (Chengdu, Chengdu herb substance CO., LTD), and ligustilide (Chengdu, Chengdu phytopurify CO., LTD).

### Plant materials

In November 2020, 50 seedlings of *L. chuanxiong* were collected from the top-geoherbalism region Pengzhou county (Sichuan, China), and ten seedlings of *L. sinense* from Lushi county (Henan, China) were collected. These seedlings were subsequently transplanted to the greenhouse of the Agricultural Genomics Institute at Shenzhen (Chinese Academy of Agricultural Sciences, China). After growing for six months, those healthy plants with similar biomass were selected for genomic sequencing, RNA-Seq, and phthalide extraction. The 4–5-week seedlings of *N. benthamiana* were used for *Agrobacterium*-mediated gene transformation. The seedlings were grown at laboratory temperature under growth lights with a 16/8 h light/dark cycle.

### Genomic DNA preparation, RNA extraction and cDNA preparation

High molecular weight genomic DNA was extracted from 3 g fresh leaves of *L. chuanxiong* stored at −80℃ using CTAB method and purified with QIAGEN® Genomic kit (QIAGEN, Germany). The DNA integrity and purity was monitored on 1% agarose gels and by NanoDropTM One UVVis spectrophotometer (Thermo Fisher Scientific, USA). DNA concentration was further measured using Qubit® 4.0 Fluorometer (Invitrogen, USA).

Total RNA was extracted from each tissue (rhizome, stem, petiole, and leaf) of *L. chuanxiong* using the RNeasy Plant Mini Kit (Omega, USA). The first-strand cDNAs were synthesized by reverse transcription PCR from the total RNA samples as templates using the SuperScript III First-Strand Synthesis System with the oligo (dT) 20 primer (Thermos Fisher Scientific, USA).

### Genome survey

The genome size of *L. chuanxiong* was estimated by flow cytometry assay. The fresh leaves of three biological replicates were chopped in LB01 lysis buffer (15 mmol/L Tris, 2 mmol/L Na^2^EDTA, 0.5 mmol/L spermine tetrahydrochloride, 80 mmol/L KCl, 20 mmol/L NaCl, 0.1% TritonX-100, 15 mmol/L β-mercaptoethano, pH 7.0–8.0) to release the nucleus. The resulting nuclei suspension was filtered through a 40 μm cell strainer, and then treated with 20 μg mL^-1^ Rnase A and 20 μg mL^-1^ propidium iodide (PI). After a 30-min incubation on ice in the dark, the fluorescence intensity of PI-stained nuclei was determined by flow cytometry (Beckman Coulter, CytoFLEX) with a sample flow rate of 10 μL/min. We used fennel (*Foeniculum vulgare*) (1C = 1.34 Gb)^77^ as the internal reference and *Zea mays* Mo17 (1C = 2.18 Gb)^78^ as the external reference. All analyses were performed in triplicate for each sample.

The genomic heterozygosity of *L. chuanxiong* was surveyed by k-mer method based on 105 Gb Illumina short-read sequencing data (∼30X). Jellyfish (v2.2.10)^79^ was performed to generate a 21-mer frequency distribution, which was then analyzed in GenomeScope 2.0^80^ to estimate heterozygosity.

### Cytogenetic and FISH assay to determine karyotype

FISH and cytogenetic assays were conducted to determine the ploidy level and chromosome number of *L. chuanxiong*. Two-centimeter root tips of seedings were cut and treated with nitrous oxide on ice for 2 h. The chromosomes were then fixed for 10 mins with 90% glacial acetic acid. After washing twice with ddH_2_O, the samples were chopped and incubated in 25 μL solution with a 3:1 mixture of 2.0% cellulase and 1.0% pectinase for 1 h at 37℃. The enzymatically hydrolyzed root tips were further mashed by tweezers and rinsed three times with 70% alcohol. The pretreated root-tip cells were suspended in 45% glacial acetic acid and dropped onto a clean slide with a cover glass at 23℃. The slides containing cells in metaphase were selected using a light microscope.

The probes (TTTAGGG) for telomere hybridization were labeled with Fluorescein-12-dUTP, a green fluorescein. Labeled probe solution of 0.25 μL mixed with 8 μL reaction buffer (1×TE and 2× SSC, pH = 7.00) was added to each slide containing cells in metaphase. The slides were incubated at 80℃ for 5 mins, and then transferred to an *in-situ* hybridization box at 40℃ for 12 h. After washing twice with ddH_2_O, the slides were stained with 8 μL DAPI and covered with cover slips. The cells in metaphase after hybridization were observed and imaged using a high-resolution fluorescence microscope (Olympus BX50, Japan).

### Genome sequencing, assembly, and quality control

For the PacBio library construction, 15 μg genomic DNA from each leaf sample of *L. chuanxiong* was fragmented to approximately 15 kb using g-TUBEs (Covaris, USA). After removing short fragments and single-strand overhangs, the retained fragments were converted into the proprietary SMRTbell library using the PacBio DNA Template Preparation Kit (Pacific Biosciences, CA, USA). Single Molecule Real Time (SMRT) sequencing was performed on a PacBio Sequel II sequencing platform.

For Hi-C library construction, chromatin was first fixed in place with formaldehyde in the nucleus and then extracted. The extracted chromatin was digested with DpnII. The 5’ overhangs of resulting fragments were then filed in with biotinylated nucleotides, and free blunt ends were ligated. After ligation, the DNA was purified from protein and treated following the Illumina Next Generation manufacturer’s instructions. The libraries were subsequently sequenced on Illumina Hiseq X, producing 296 Gb 2 × 150 bp paired-end reads.

The raw data of PacBio subreads was filtered to HiFi reads by PBccs (v6.4.0) (https://github.com/PacificBiosciences/ccs), and subsequently assembled using Hifiasm (v0.16.0)^81, 82^. The initial assembled contigs were anchoring to chromosomes by 3D-DNA pipeline (v201008)^83^ and further manual adjustments were made to produce a chromosome-level genome. BUSCO (v5.4.3)^84^ was used for benchmarking the genome with “embryophyte_odb10” database.

### Genome annotation

EDTA (v2.0.0)^85^ was used to *de novo* identify, annotate, and classify the repetitive elements in the genome of *L. chuanxiong*. Before protein-coding gene annotation, the annotated repetitive elements in genome were soft masked using bedtools (v2.28.0)^86^. RNA-Seq raw reads of *L. chuanxiong* were filtered using fastx-toolkits (v0.0.14) (http://hannonlab.cshl.edu/fastx_toolkit/index.html) and then assembled through Hisat2 (v2.2.1)^87^ and Stringtie (v2.2.0)^88, 89^. The raw assembly of transcripts were further validated by PASA (v2.5.1)^90^, which were then incorporated to the MAKER (v3.01.03)^91^ pipeline to automatedly identify protein-coding genes. Finally, the gene models identified by MAKER (v3.01.03)^91^ were updated by PASA (v2.5.1)^90^. Functional annotations of the protein-coding genes were carried out by using BLASTP searches against entries in both NCBI non-redundant protein (NR) (https://www.ncbi.nlm.nih.gov/) and Swiss-Prot (https://www.uniprot.org/) databases. The prediction of conserved domains for the genes was performed by InterProScan (v5.11-51.0)^92^. The annotations of the GO terms (http://geneontology.org/) and KEGG pathways (https://www.genome.jp/kegg/) for the genes were added using eggNOG-mapper (v2.1.10-0)^93^. Circos plot was visualized using TBtools^94^.

### Haplotype phasing for the genome

SubPhaser (v1.2)^43^ pipeline was used to perform the genome phasing analysis. Jellyfish (v2.2.10)^79^ was firstly performed to generate a 17-mer frequency distribution. Then a K-Means algorithm was employed to cluster k-mers throughout the whole genome. The significance level of k-mers clusters was evaluated by bootstrap method. Finally, subgroups within genome were identified by performing hierarchical clustering and principal component analysis (PCA).

### Detection of SVs and SNPs

Given the large genetic distance between the two haplotypes of *L. chuanxiong*, it was challenging to accurately and directly aligned the whole genomic sequence of *Lc*HapA and *Lc*HapB. Thus, we used the syntenic gene block strategy to align and detect the SVs between haplotypes. Jcvi (v1.1.18)^95^ was used to detect collinear gene blocks between *Lc*HapA and *Lc*HapB, and only those unique gene block pairs (1:1) were remained for SV detection. Possible duplications caused by ancient WGD events or other fragment duplication events were removed to avoid overestimate of SVs.

In order to compare the sequence similarity among *Lc*HapA, *Lc*HapB and *L. sinense*, we performed SNP calling for *Lc*HapA and *L. sinense* using *Lc*HapB genome as the reference based on RNA-Seq data. First, we performed RNA-Seq for *L. sinense*. Second, we extracted the *Lc*HapA specific reads from the total reads initially mapped to a combined reference of two haplotypes, using the *Lc*HapA genome. Third, RNA-Seq reads of *Lc*HapA and *L. sinense* were mapped to *Lc*HapB using Hisat2 (v2.2.1)^87^. Subsequently, GATK (v4.2.4.0)^96^ best practice workflow was employed to call SNPs with parameter “--sample-ploidy 1 --intervals”. Finally, the SNP number scanned by 100-kb non-overlapped sliding windows was compared between *Lc*HapA and *L. sinense*.

### *De novo* assembly of transcriptome of *L. sinense*

RNA-Seq raw reads of *L. sinense* were filtered using fastx-toolkits (v0.0.14) (http://hannonlab.cshl.edu/fastx_toolkit/index.html) and then *de novo* assembled into transcripts using Trinity (v2.13.2)^97^. The longest transcript was chosen by Trinity in-house scripts. TransDecoder (v5.5.0) (https://github.com/TransDecoder/TransDecoder) was used to predict CDS. The orthologous gene sets among *Lc*HapA, *Lc*HapB and *L. sinense* were obtained using the bi-directional best matching (BBH) method performed through BLAST (v2.13.0+)^98^. Finally, the CDS identity of each orthologous gene pair among *Lc*HapA, *Lc*HapB and *L. sinense* was calculated by MEGA11^99^.

### Comparative genomic analyses

In order to clarify the phylogenetic relationship and divergence time among *Lc*HapA, *Lc*HapB, *L. sinense* and the other 10 species (**Fig. 3d**; **Supplementary Table 7**), we constructed ML trees based on single-copy orthologous genes obtained from RNA-Seq data. In total, we identified 193 single-copy orthologs using OrthoFinder (v2.5.4)^100^ with default settings, and aligned each of them by MUSCLE (v5.1.linux64)^101^. These aligned orthologs were concatenated by Phylosuite (v1.2.2)^102^. IQTree2 (v2.2.0.3)^103^ was used to detect the best-fit amino acid substitution model, based on which RAxML-NG (v1.1.0)^104^ was used to construct the ML phylogeny with 1,000 bootstrap analyses. MCMCTree tool of PAML package (v4.9j)^105^ was used to estimate the divergence time, with three constraints for calibration^106^ (**Fig. 3d**; **Supplementary Table 15**).

WGDI (v0.6.1)^107^ was used to detect the WGD events in Apiales. The gene family expansion and contraction analyses were performed using CAFE5 (v5.0)^108^, where the gamma rate was set from 1 to 7, and “k = 5” was selected for best fit. GO enrichments were performed by ClusterProfiler (v4.6.0)^109^ for those significantly expanded or contracted gene families. The results of analyses were visualized using R package ggtree (v3.6.2)^110^.

### Allele-specific expression (ASE) analyses

To determine the ASEs, we conducted comprehensive comparison of gene expression level between *Lc*HapA and *Lc*HapB in different genomic structure (linear, inversion, and translocation regions) and tissue (rhizome, leaf, petiole, stem) types. We initially identified allele pairs in each syntenic gene block based on BBH methods by BLAST (v2.13.0+)^98^. Then, we quantified the expression level of each allele based on RNA-Seq data. Clean RNA-Seq reads were mapped to *Lc*HapA and *Lc*HapB through Hisat2 (v2.2.1)^87^ and assembled by Stringtie (v2.2.0) ^88, 89^. Expression count matrix for all allele pairs was constructed by a script prepDE.py downloaded from the stringtie website (http://ccb.jhu.edu/software/stringtie/dl/prepDE.py). Based on this matrix, ASEGs were identified using DESeq2^111^ (*P* < 0.05 and FDR < 0.05, log2(foldchange) > 1). *K*a and *K*s of all allele pairs were calculated by *K*s module of WGDI (v0.6.1)^107^. Student’s t test was used for comparisons of *K*a, *K*s, or *K*a/*K*s between ASEGs and non-ASEGs in each genomic structure and tissue type with a significance level of *P* < 0.05. Fisher’s exact test was used to compare the ASEGs frequency between SV and non-SV regions with a significance level of *P* < 0.05.

### Phthalide extraction and GC-MS analysis

Approximately 100 mg of tissue samples were dissected and homogenized to a fine powder on a ball mill. Phthalides were extracted using 5–10 volumes (w/v) of ethyl acetate at room temperature for 1 h. The extracts were centrifuged twice (13,000 g, 20 min) and supernatants were collected for GC–MS analysis. The conditions of GC-MS analyses for phthalide mainly followed other studies^112, 113^.

GC-MS analysis was carried out on an Agilent 7890B GC machine (Agilent Technologies, Waldbronn, USA) with an Agilent 7000C mass selective detector at 70 eV and a helium flow of 1.0 mL min^-1^. The sample of 1 µL was injected at a split ratio of 50:1 and analyzed on an Agilent HP-5MS column. The column temperature was set at 50℃ for injection, then programmed at 5℃/min to 250℃. The spectrometers were operated in electron-impact (EI) mode and the phthalides were analyzed with the selected reaction monitoring (SRM) mode. The inlet and ionization source temperatures were set at 280℃ and 300℃, respectively. The mass spectrum of l-NBP, butylidenephthalide, senkyunolide A, and ligustilide were presented in **Supplementary Fig. 27**.

### Crude protein feeding assay

The crude protein feeding assay mainly followed the protocol of Vanholme et al. (2013)^114^. Crude protein from ∼200 mg of ground rhizome tissue of *L. chuanxiong* was extracted by 1 mL protein extraction buffer on ice for 1 h (inverting the tubes every 5–10 mins). We then centrifuged the extract at 14,000 rpm for 10 min at 4°C and transferred supernatant to a new precooled tube. Protein concentration of the rhizome crude protein extract (CPE) was quantified by pierce 660 nm protein assay reagent. At last, we fed the CPE with 40 μg of each target phthalide (l-NBP, butylidenephthalide, senkyunolide A, and ligustilide) and subsequently detect signals for the other three by GC-MS.

### Genome wide mining for P4,5Ds

We initially excluded those genes with a FPKM ≤ 20 and selected those significantly up-regulated genes in the rhizome (log2(foldchange) > 1, *P* value < 0.05, FDR < 0.05) compared with leaf, stem, and petiole using DESeq2^111^. We then calculated Pearson correlation coefficients between contents of l-NBP (also butylidenephthalide) and the FPKM of genes in rhizome using the R package psych (https://rdocumentation.org/packages/psych/versions/2.3.3), and selected those genes with r > 0.95 and *P* < 0.05.

HMMER3 (v3.3.2)^115^ was used to search 2OGDs and CYPs with an e-value of 1e^-6^. HMMER profile PF00067 was used for CYPs search, and PF14226 and PF03171 were used to search genes in DOXC family (2OGD superfamily) that was usually involved in plant specificized metabolite pathway^55^. The possible pseudogenes (length of predicted CYPs < 300 amino acids and 2OGDs < 200 amino acids) were discarded. Gene structures of all candidate 2OGDs and CYPs were manually adjusted using IGV-GSAman (https://gitee.com/CJchen/IGV-sRNA).

At last, we constructed ML phylogenies using RAxML-NG (v1.1.0)^104^ for candidate CYPs and 2OGDs. The protein alignment was obtained using MUSCLE (v5.1.linux64)^101^. The best fitting amino acid substitution model was selected as LG+I+G4+F for both CYPs and 2OGDs by ModelTest-NG (v0.1.7)^116^. For each 2OGD (DOXC family) clade or CYP family, we only selected 1–3 highly-expressed genes for cloning.

### Cloning of candidate P4,5Ds

All biosynthetic genes tested in the present study were cloned from the cDNA of *L. chuanxiong*. CDS for each gene was amplified via PCR using KOD One^TM^ PCR master Mix (Toyobo). Oligonucleotide primers were designed to target the 5’ and 3’ homologous regions for subsequent homologous recombination assembly into the appropriate expression plasmid. Amplicon products of PCR were analyzed by gel electrophoresis on a 1% (w/v) agarose gel, and the positive bands were excised and purified using the Omega Gel DNA Recovery Kit (Omega Research). A list of all primers used for CDS cloning in this study was provided in **Supplementary Table 16**.

### Functional characterization for candidate 2OGDs in *E. coil* system

Candidate 2OGDs were cloned into the pMAl vector, a bacterial expression vector based on maltose binding protein tag (MBP), following the manufacture’s handbook. The purified plasmids were transformed into Rosetta (DE3) strains of *E. coli* using heat shock transformation. Transformed cells were selected overnight on Lysogeny broth (LB) plates with ampicillin at 37°C. The positive colonies were screened by colony PCR and further cultured in 100 mL LB containing ampicillin at 200 rpm and 37°C until the OD600 absorbance reaching 0.5–0.6, when 0.5 Mm isopropyl-β-D-thiogalactopyranoside (IPTG) (MedChemExpress) was added to induce protein expression. At the same time, 50 μM senkyunolide A or ligustilide was added. The strains were further cultured at 37°C for 4 h under 200 rpm. At last, the product was extracted with 3-fold volume of ethyl acetate and then analyzed by GC-MS.

### Functional characterization for candidate CYPs in *N. benthamiana* system

Candidate CYPs were cloned into the pSuper1300 vector and transformed into the GV3101 strains of *Agrobacterium* electrocompetent cells. The strains were cultured overnight in LB medium with 50 µg/mL kanamycin at 28℃, 180 rpm. Then, 500 µL of *Agrobacterium* suspension was transferred into 30 mL fresh LB medium containing rifampicin (25 µg/mL) and kanamycin (50 µg/mL) and cultured overnight. *Agrobacterium* cells were collected by centrifugation at 4,000 rpm for 10 min and resuspended with induction medium (100 µM acetosyringone, 10 mM MES, 10 mM MgCl_2_, pH = 5.6) for 1–2 hours at room temperature. The OD600 absorbance of each strain resuspension was determined and diluted with induction medium to ∼0.3. Each diluted strain was then infiltrated into the abaxial side of three leaves from different 4–5-week *N. benthamiana* seedlings using a 1 mL needleless syringe.

To test the enzymatic activity, senkyunolide A or ligustilide (not present in *N. benthamiana*) solution (100 μM substrate in DI water) was co-infiltrated into the *Agrobacterium*-infiltrated leaves after 3 days. The infiltrated leaf region with substrate was marked and subsequently taken for phthalide extraction after one day. Infiltrations for all treatments regarding different genes and substrates were conducted in triplicate.

## Data availability

The raw data of genome and transcriptome sequencing of *L. chuanxiong* and *L. sinense* have been deposited to the Genome Sequence Archive at the National Genomics Data Center (NGDC) under BioProject No. PRJCA017485. For review purposes, *L. chuanxiong* genome files are available at figshare [https://figshare.com/s/21f94c7dfbc48759aef3].

## Supporting information

The supplementary files contain all supplementary figures and supplementary tables cited in the main text.

## Acknowledgements

This study was funded by National Key Research and Development Program of China (Grant No. 2020YFA0907900), National Natural Science Foundation of China (Grant No. 32070242), Shenzhen Science and Technology Program (Grant No. KQTD2016113010482651), Special funds for Science Technology Innovation and Industrial Development of Shenzhen Dapeng New District (Grant Nos. RC201901-05 and PT201901-19), China Postdoctoral Science Foundation (Grant No. 2020M672904), Basic and Applied Basic Research Fund of Guangdong (Grant No. 2020A1515110912), and Science, Technology and Innovation Commission of Shenzhen Municipality of China (ZDSYS 20200811142605017). We would like to thank Zhaolong Duan for providing seedings of *L. chuanxiong* and Zengxiang Guo for providing seedlings of *L. sinense*. We thank Jindan Zhang for providing suggestions on experimental details of cytogenetic assay. We thank Yanchun Peng for assistance in GC-MS analysis. We thank Zhigui Bao for his help in the early stages of this project. We thank Xianjin Qin and Dejin Xie for their help in the *in vitro* enzyme activity assay. We also thank David Nelson of P450 nomenclature committee for naming the CYPs.

## Author contributions

L.W., X.C., Z.H., and B.N. conceived and designed the study. B.N. and L.Z. prepared the plant materials for sequencing. Z.H., B.N., and C.L. conducted the bioinformatics analyses. B.N. conducted the FISH analysis. J.J. and L.Z. performed the cytogenetic assay. X.C. and W.S. cloned and characterized the candidate genes. X.C., W.S, and S.Y. performed GC-MS analyses. B.N., X.C., and Z.H. developed the figures, interpreted the results, and wrote the manuscript. L.W., W.L., C.L., H.L., Y.T., W.Z., and P.F. provided suggestions on manuscript revising. All authors read and approved the final version of the manuscript.

## Competing interests

The authors declare no competing interests.

## Notes

### Competing Interest Statement

The authors have declared no competing interest.

