## Supplementary material for "Haplotype-phased genome revealed the butylphthalide biosynthesis and hybrid origin of *Ligusticum chuanxiong*": The supplementary files contain all supplementary figures and supplementary tables cited in the main text.: Chuanxiong_NBP_SuppInfo_NC_clean.pdf

### SUPPLEMENTARY INFORMATION

**Supplementary Figure 1.** Hypotheses for the biosynthesis pathway of phthalide scaffold in *L. chuanxiong*.

**Supplementary Figure 2.** Mono-phthalide diversity in *L. chuanxiong*.

**Supplementary Figure 3.** C-value and karyotype of *L. chuanxiong*.

**Supplementary Figure 4.** Intensity heat map of Hi-C chromosome interaction for *L. chuanxiong* genome, with a resolution of 2.5 Mb.

**Supplementary Figure 5.** Haplotype phasing of *L. chuanxiong* genome.

**Supplementary Figure 6.** Gene family expansion and contraction patterns tracing along the phylogeny of Apiales and the number of annotated genes of each species.

**Supplementary Figure 7.** GO enrichment for the expanded gene families along the lineage of *Lc*HapA (a), *Lc*HapB (b), the most recent ancestor of *Lc*HapA and *Lc*HapB (c), and the crown node of *L. chuanxiong* and *Angelica sinensis* (d).

**Supplementary Figure 8.** GO enrichment for the contracted gene families along the lineage of *Lc*HapA (a), *Lc*HapB (b), the most recent ancestor of *Lc*HapA and *Lc*HapB (c), and the crown node of *L. chuanxiong* and *A. sinensis* (d).

**Supplementary Figure 9.** Identification of WGD events in Apiales.

**Supplementary Figure 10.** Synteny among *Lc*HapA, *Lc*HapB, and *A. sinensis*.

**Supplementary Figure 11.** The inter-haplotype or inter-subgenome SVs for several diploid and alloploid species.

**Supplementary Figure 12.** Mapping rate and mapping quality of the RNA-Seq reads of *L. sinense* onto genomes of *Lc*HapA and *Lc*HapB.

**Supplementary Figure 13.** Summary of the possible phylogenetic topologies among *Lc*HapA, *Lc*HapB and other species.

**Supplementary Figure 14.** The distribution of ASEGs on different chromosomes and among tissues.

**Supplementary Figure 15.** The comparisons for *K*s and *K*a/*K*s ratio of ASEGs and non-ASEGs in *L. chuanxiong* and the GO enrichment of ASEGs.

**Supplementary Figure 16.** The Venn diagram of the number of significantly up-regulated genes with high expression (FPKM > 20) in rhizome compared to the other three tissues of *L. chuanxiong*.

**Supplementary Figure 17.** The ML phylogenetic tree of candidate 2OGD (DOXC family) genes for l-NBP (**1**) and butylidenephthalide (**2**) biosynthesis in *L. chuanxiong*.

**Supplementary Figure 18.** The ML phylogenetic tree of candidate CYPs for l-NBP (**1**) and butylidenephthalide (**2**) biosynthesis in *L. chuanxiong*.

**Supplementary Figure 19.** GC-MS peak plots of l-NBP (**1**) (*m/z* 133) in the indicated combinations of senkyunolide A (**3**) and candidate 2OGDs.

**Supplementary Figure 20.** GC-MS peak plots of butylidenephthalide (**2**) (*m/z* 159) in the indicated combinations of ligustilide (**4**) and candidate CYPs.

**Supplementary Figure 21.** The ML phylogeny of characterized plant 2OGD (DOXC family).

**Supplementary Figure 22.** The ML phylogeny of characterized plant CYP716s.

**Supplementary Figure 23.** The ML phylogeny of characterized plant CYP72s.

**Supplementary Figure 24.** The tandem duplication of CYP716Es in Apiaceae.

**Supplementary Figure 25.** The ML phylogeny of Apiaceae specific tandem duplicated CYP716Es.

**Supplementary Figure 26.** Comparison of the content of l-NBP between *L. chuanxiong* and *L. sinense*.

**Supplementary Figure 27.** The mass spectrum of l-NBP (**1**), butylidenephthalide (**2**), senkyunolide A (**3**), and ligustilide (**4**).

**Supplementary Table 1.** Summary of the sequencing data of *L. chuanxiong* generated in the present study.

**Supplementary Table 2.** Summary of the haplotype-phased genome of *L. chuanxiong*.

**Supplementary Table 3.** Summary of the annotation features of each chromosome of *L. chuanxiong*.

**Supplementary Table 4.** Repetitive elements annotation in the genome of *L. chuanxiong*.

**Supplementary Table 5.** Summary of the SVs between *Lc*HapA and *Lc*HapB.

**Supplementary Table 6.** Summary of the RNA-Seq assembly of *L. sinense*.

**Supplementary Table 7.** The genome data sources of other species used in the present study.

**Supplementary Table 8.** Comparisons for ASEs between SV and non-SV regions in *L. chuanxiong*.

**Supplementary Table 9.** Comparisons for *Lc*HapA-bias genes between SV and linear regions in *L. chuanxiong*.

**Supplementary Table 10.** Comparisons for *Lc*HapB-bias genes between SV and linear regions in *L. chuanxiong*.

**Supplementary Table 11.** Candidate 2OGDs and CYPs for phthalide C-4,5 desaturations.

**Supplementary Table 12.** The characterized plant 2OGDs (DOXC family).

**Supplementary Table 13.** The characterized plant CYP72s.

**Supplementary Table 14.** The characterized plant CYP716s.

**Supplementary Table 15.** Calibration constraints used for divergence time estimation.

**Supplementary Table 16.** Primers used for gene cloning in the present study.

#### Supplementary Figure:

**Scheme 1**

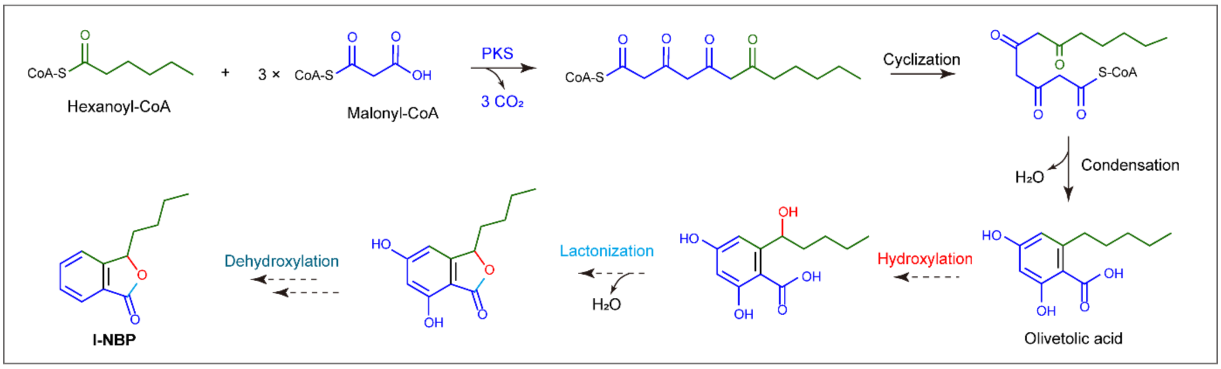


Polyketide pathway

**Scheme 2**

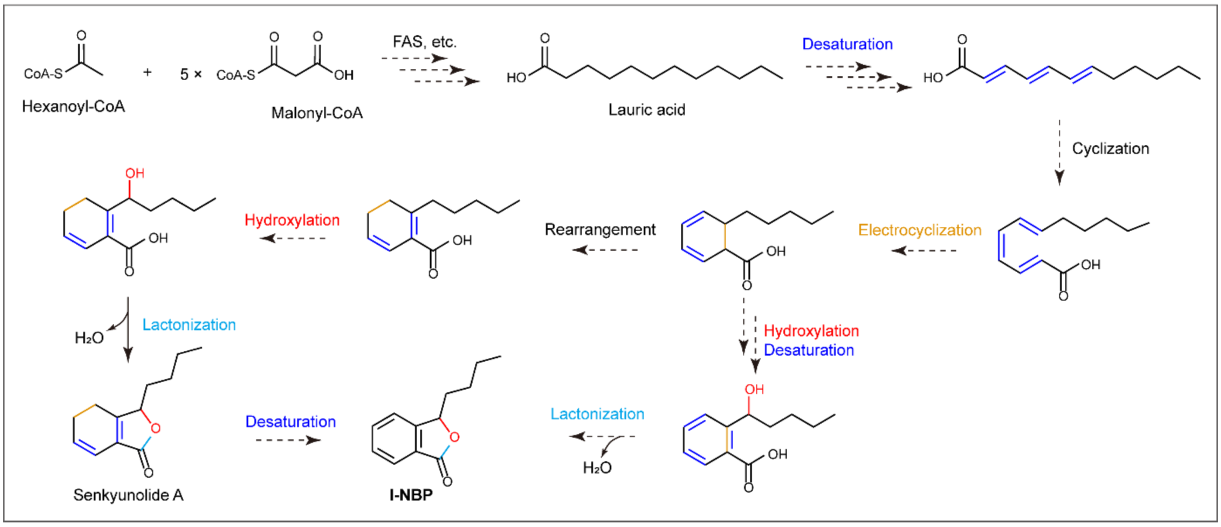


Fatty acid pathway

**Supplementary Figure 1. Hypotheses for the biosynthesis pathway of phthalide scaffold in *L. chuanxiong*.** Scheme 1 proposed a polyketide pathway, where the precursor was olivetolic acid whose biosynthesis was previously examined in *Cannabis sativa*[1,2] and *Helichrysum umbraculigerum*[3]. Scheme 2 presented a fatty acid pathway where lauric acid or its unsaturated derivative containing 12 carbons was hypothesized as the precursor for phthalide biosynthesis. The solid lines represent characterized steps in *C. sativa*[1,2], while the dotted lines represent possible but unresolved steps.

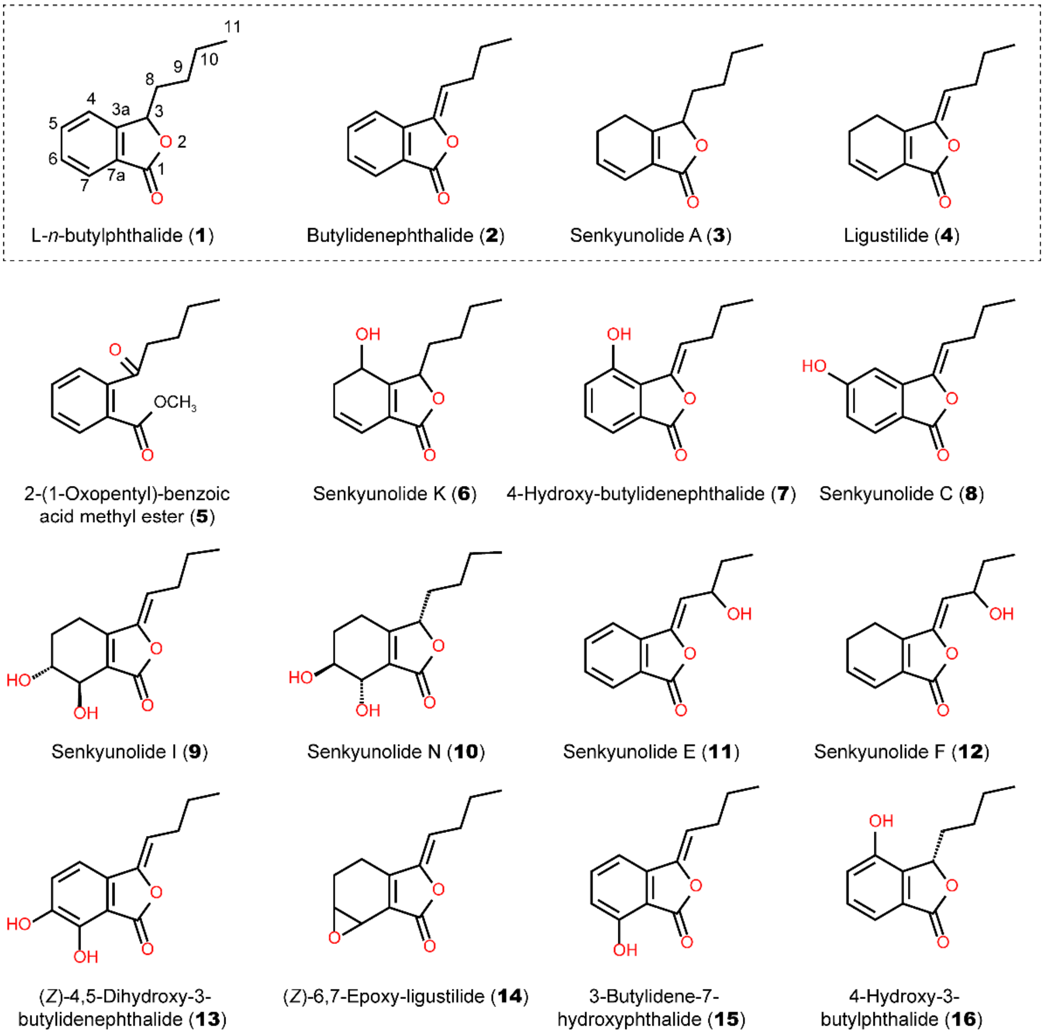


**Supplementary Figure 2. Mono-phthalide diversity in *L. chuanxiong*.** Chemicals **1–16** are examples of phthalides isolated from *L. chuanxiong*[4,5], with the four main mono-phthalides highlighted within a dashed rectangle.

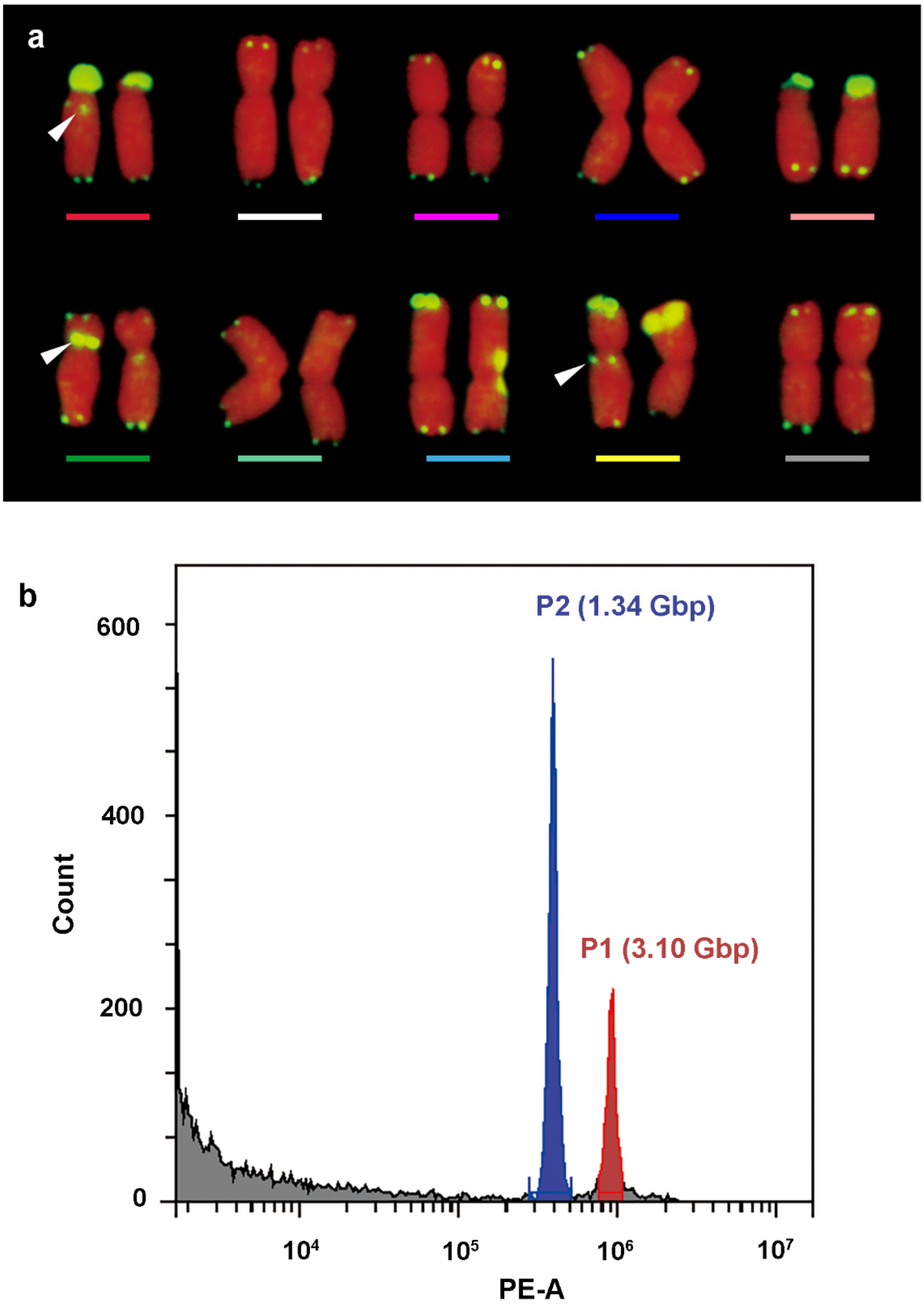


**Supplementary Figure 3. C-value and karyotype of *L. chuanxiong*. a.** Chromosome number and telomeric repeats distribution (green) in each chromosome detected by FISH assay. The white arrows indicate asymmetric signals between homologous chromosomes. **b.** C-value of *L. chuanxiong* (P1 in red) determined by flow cytometry with *Foeniculum vulgare* (P2 in blue) as an internal reference.

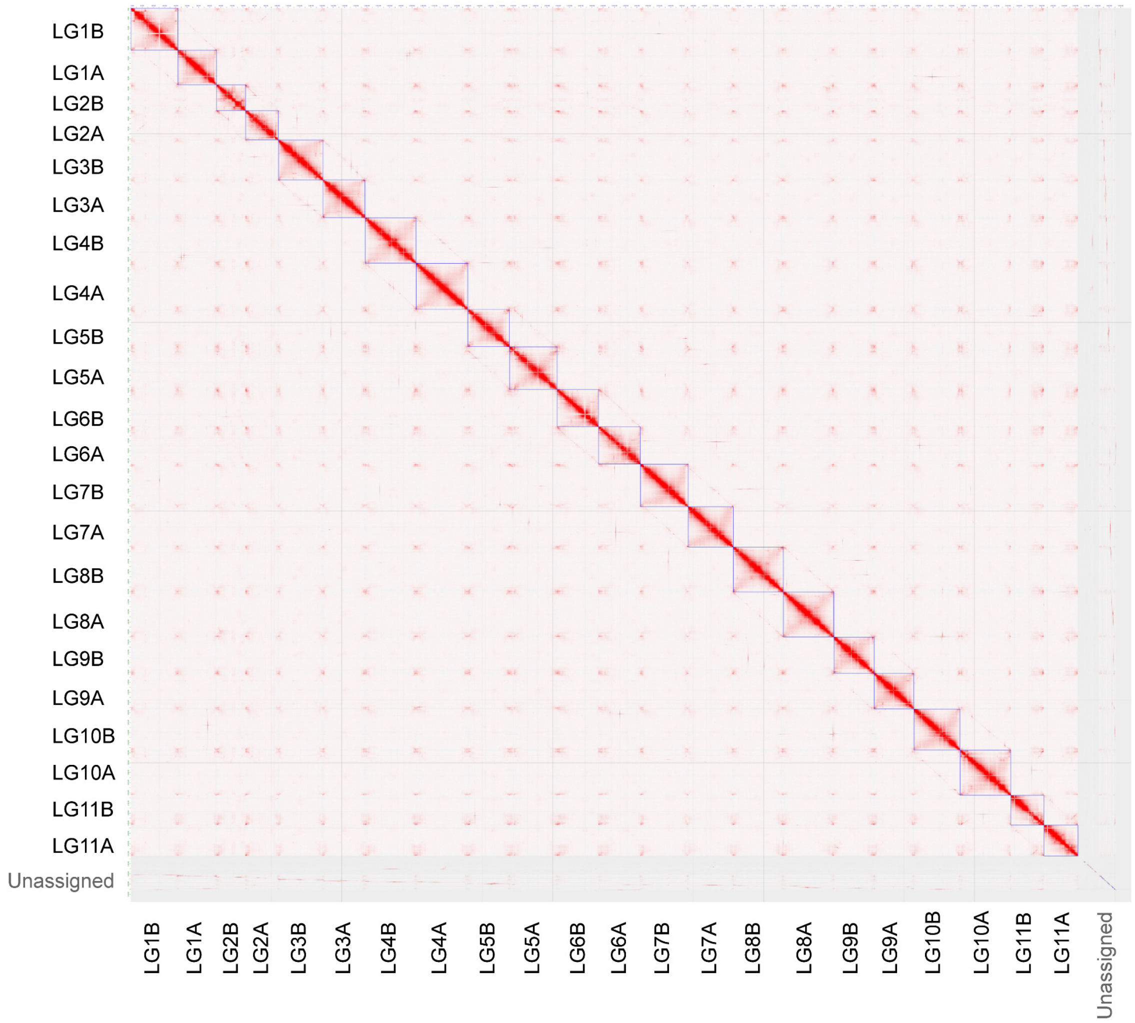


**Supplementary Figure 4. Intensity heat map of Hi-C chromosome interaction for *L. chuanxiong* genome, with a resolution of 2.5 Mb.**

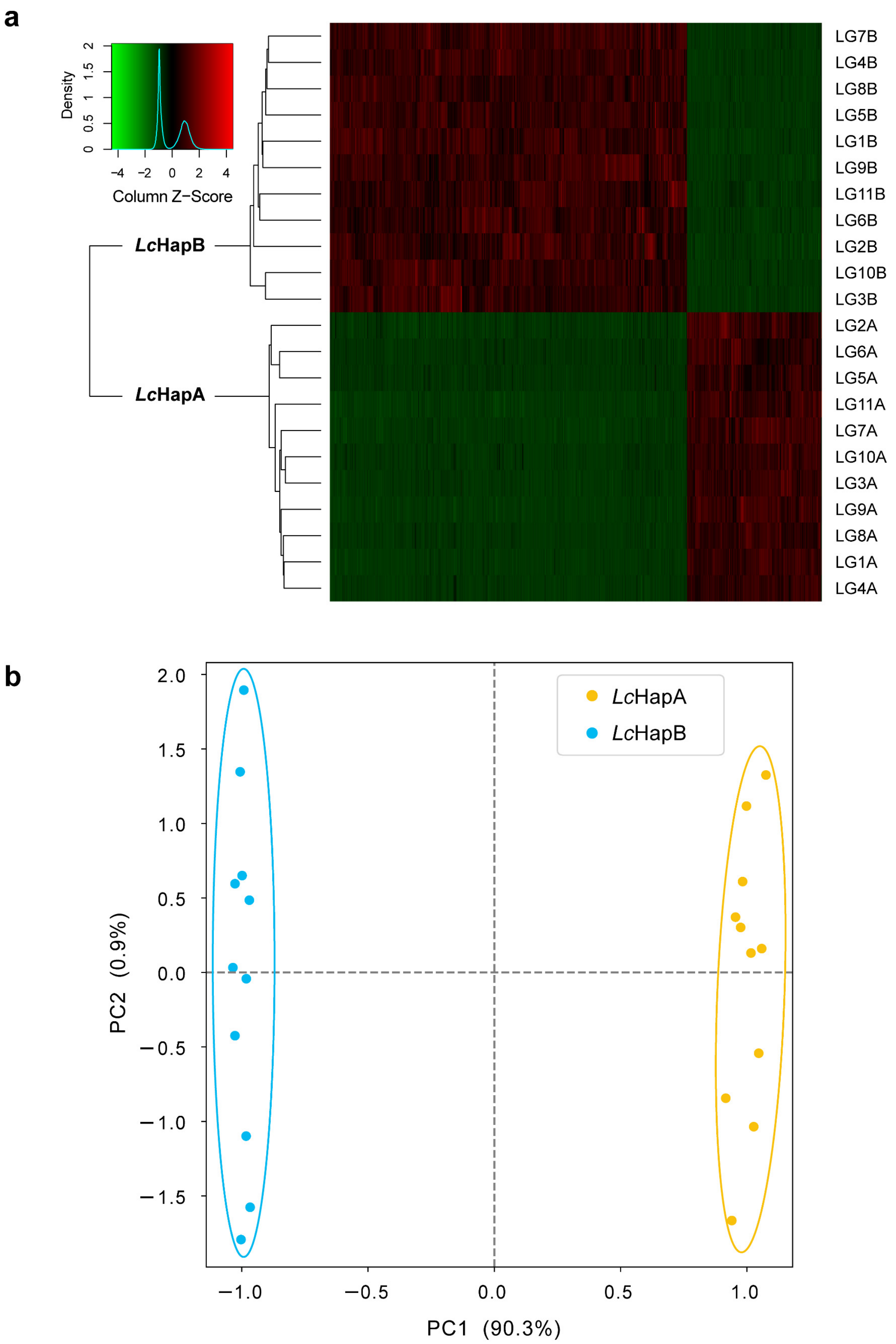


**Supplementary Figure 5. Haplotype phasing of *L. chuanxiong* genome. a.** Hierarchical clustering for 22 chromosomes based on haplotype specific kmers by SubPhaser[6]. **b.** PCA analysis for 22 chromosomes based on haplotype specific kmers by SubPhaser[6].

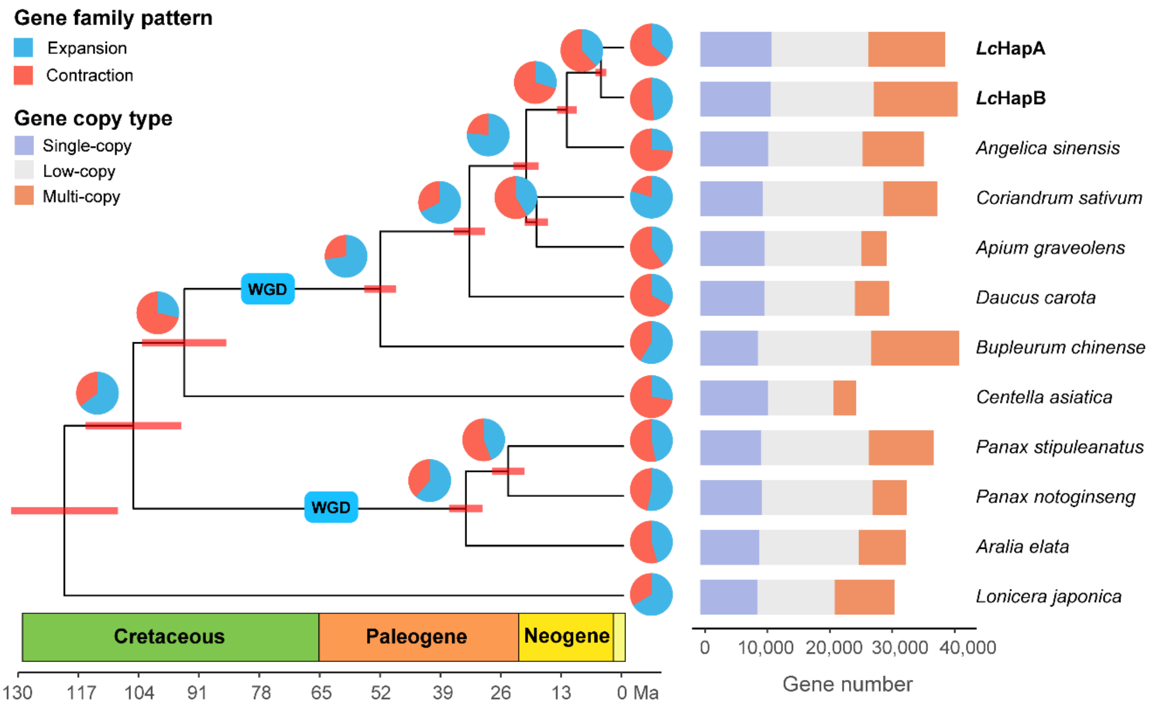


**Supplementary Figure 6. Gene family expansion and contraction patterns tracing along the phylogeny of Apiales and the number of annotated genes of each species.** The WGD events were labeled in blue rectangles. Low-copy gene, gene with a copy number of 2–5; multi-copy gene, gene with a copy number higher than five.

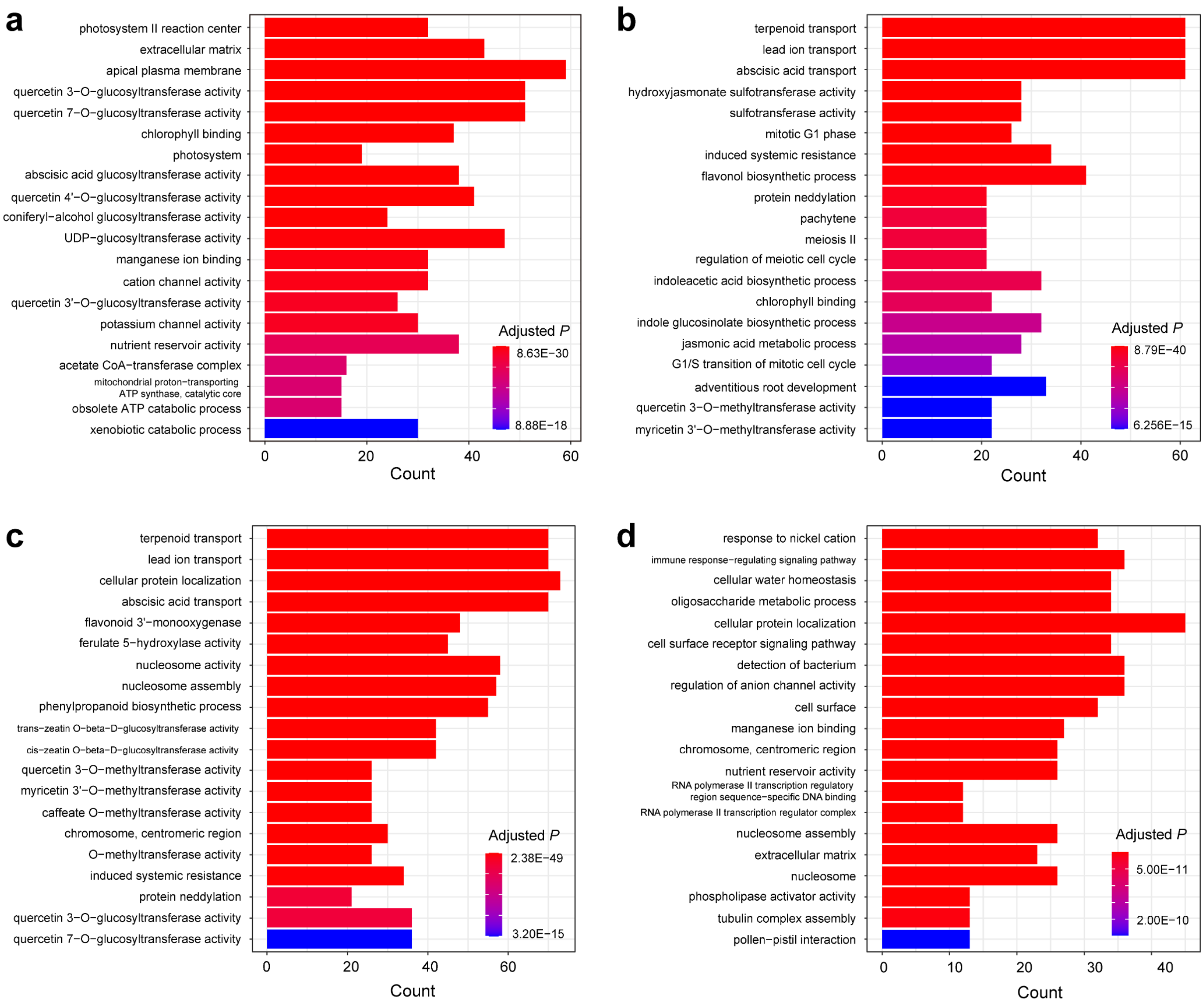


**Supplementary Figure 7. GO enrichment for the expanded gene families along the lineage of *Lc*HapA (a), *Lc*HapB (b), the most recent ancestor of *Lc*HapA and *Lc*HapB (c), and the crown node of *L. chuanxiong* and *Angelica sinensis* (d).**

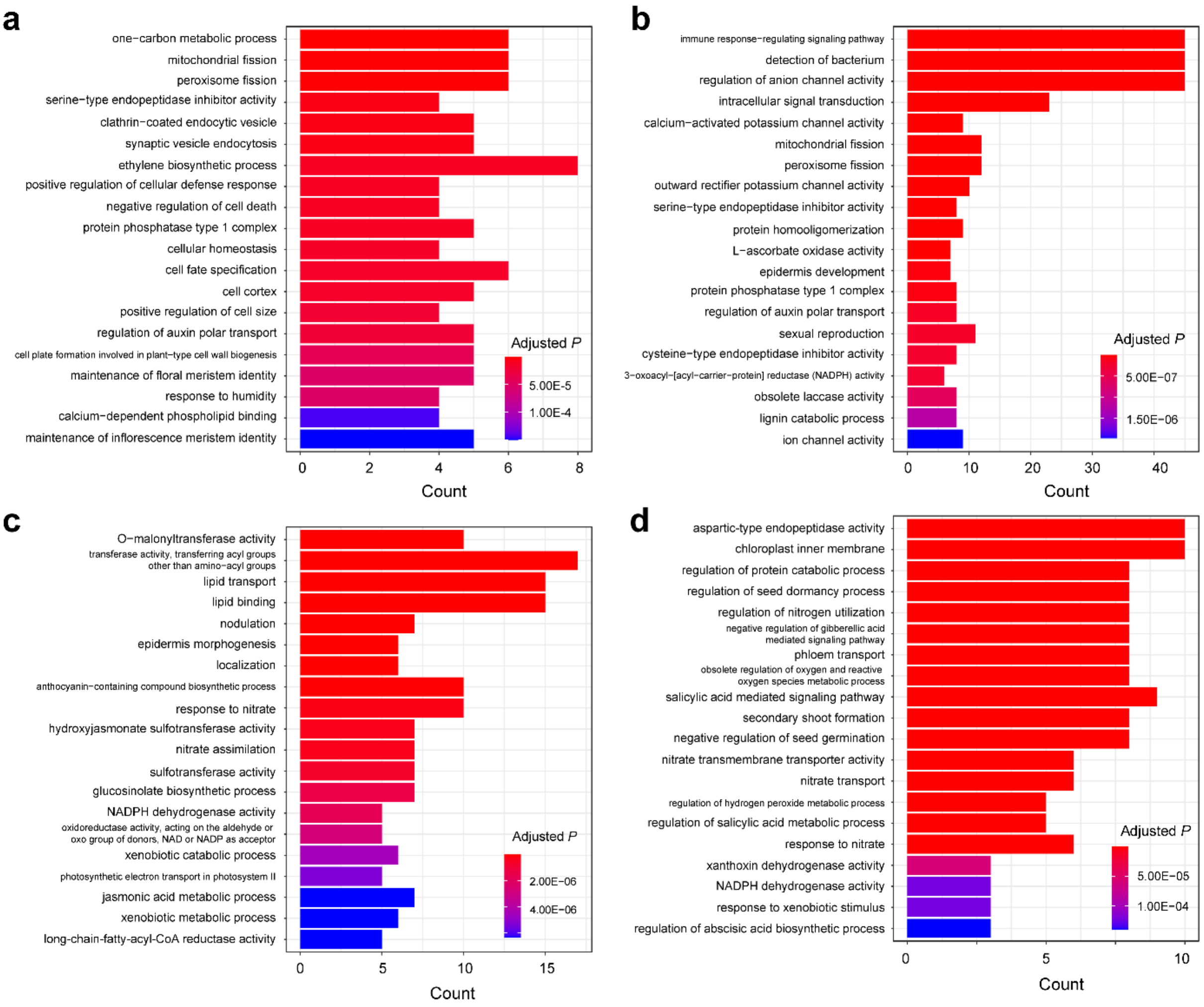


**Supplementary Figure 8. GO enrichment for the contracted gene families along the lineage of *Lc*HapA (a), *Lc*HapB (b), the most recent ancestor of *Lc*HapA and *Lc*HapB (c), and the crown node of *L. chuanxiong* and *A. sinensis* (d).**

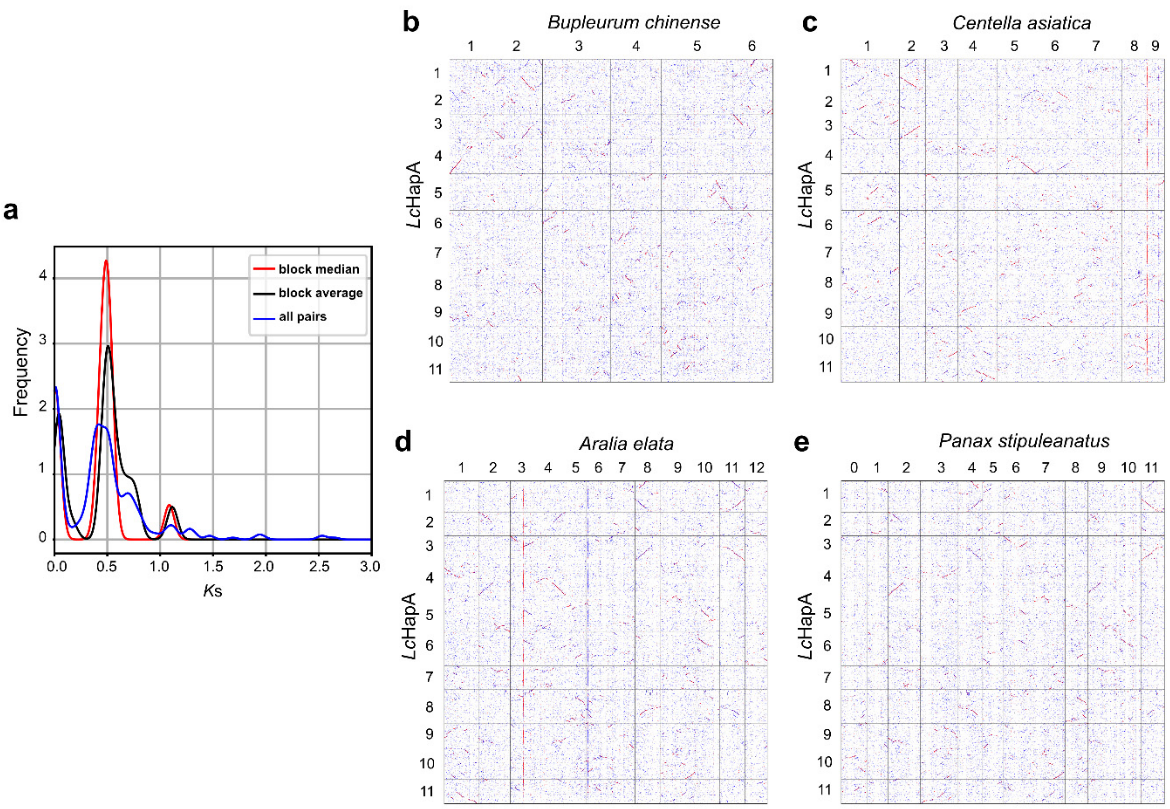


**Supplementary Figure 9. Identification of WGD events in Apiales. a.** Two *K*s peaks were detected in *L. chuanxiong*, which represented two WGD events shared by core eudicots (WGD-γ) and Apioideae. **b.** Roughly 1:1 syntenic depth ratio was detected between *Lc*HapA and *B. chinense* (one basal species of Apioideae subfamily), indicating the WGD event was shared by all species of Apioideae subfamily. **c.** Roughly 2:1 syntenic depth ratio was detected between *Lc*HapA and *C. asiatica* (belonging to the basal subfamily of the Apiaceae, Mackinlayoideae), indicating the WGD event may be specific to Apioideae. **d–e.** Roughly 2:2 syntenic depth ratio was detected between *Lc*HapA and *Aralia elata* (Araliaceae), as well as between *Lc*HapA and *Panax stipuleanatus* (Araliaceae), confirming only one WGD event in both Apiaceae and Araliaceae. The single Araliaceae-common WGD event has been reported by Wang et al. (2022)[7].

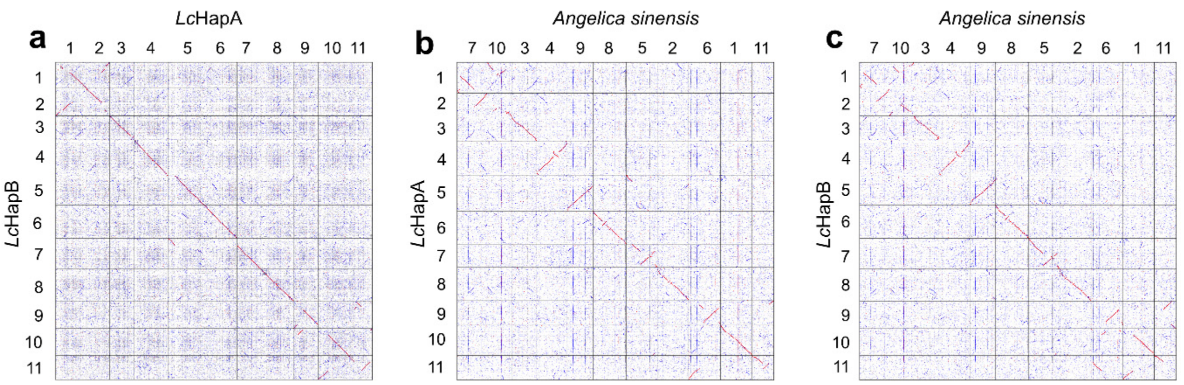


**Supplementary Figure 10. Synteny among *Lc*HapA, *Lc*HapB, and *A. sinensis*. a.** Remarkable chromosome structural differences were detected between *Lc*HapA and *Lc*HapB in Chr1, Chr2, Chr5, Chr9, Chr10, and Chr11. Synteny between *A. sinensis* and *Lc*HapA **(b)** as well as *Lc*HapB **(c)**. The Chr 1 of *A. sinensis* exhibited better synteny with Chr10 of *Lc*HapA than *Lc*HapB.

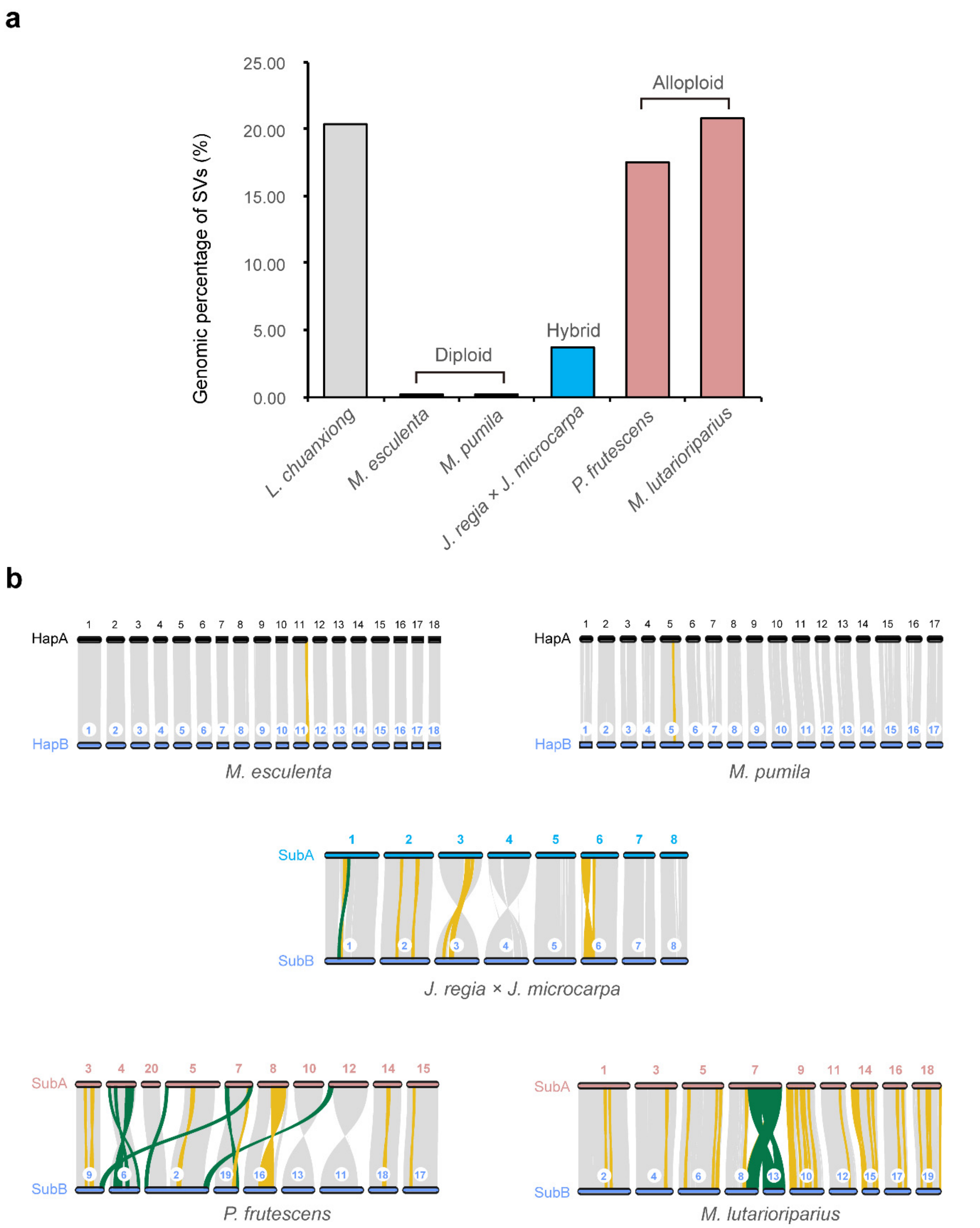


**Supplementary Figure 11. The inter-haplotype or inter-subgenome SVs for several diploid and alloploid species. a.** The genomic percentage of haplotypic or subgenomic SVs in six species. **b.** Inter-haplotype or inter-subgenome synteny of *M. esculenta*, *M. pumila*, *J. regia* × *J. macrocarpa*, *P. frutescens*, and *M. lutarioriparius*.

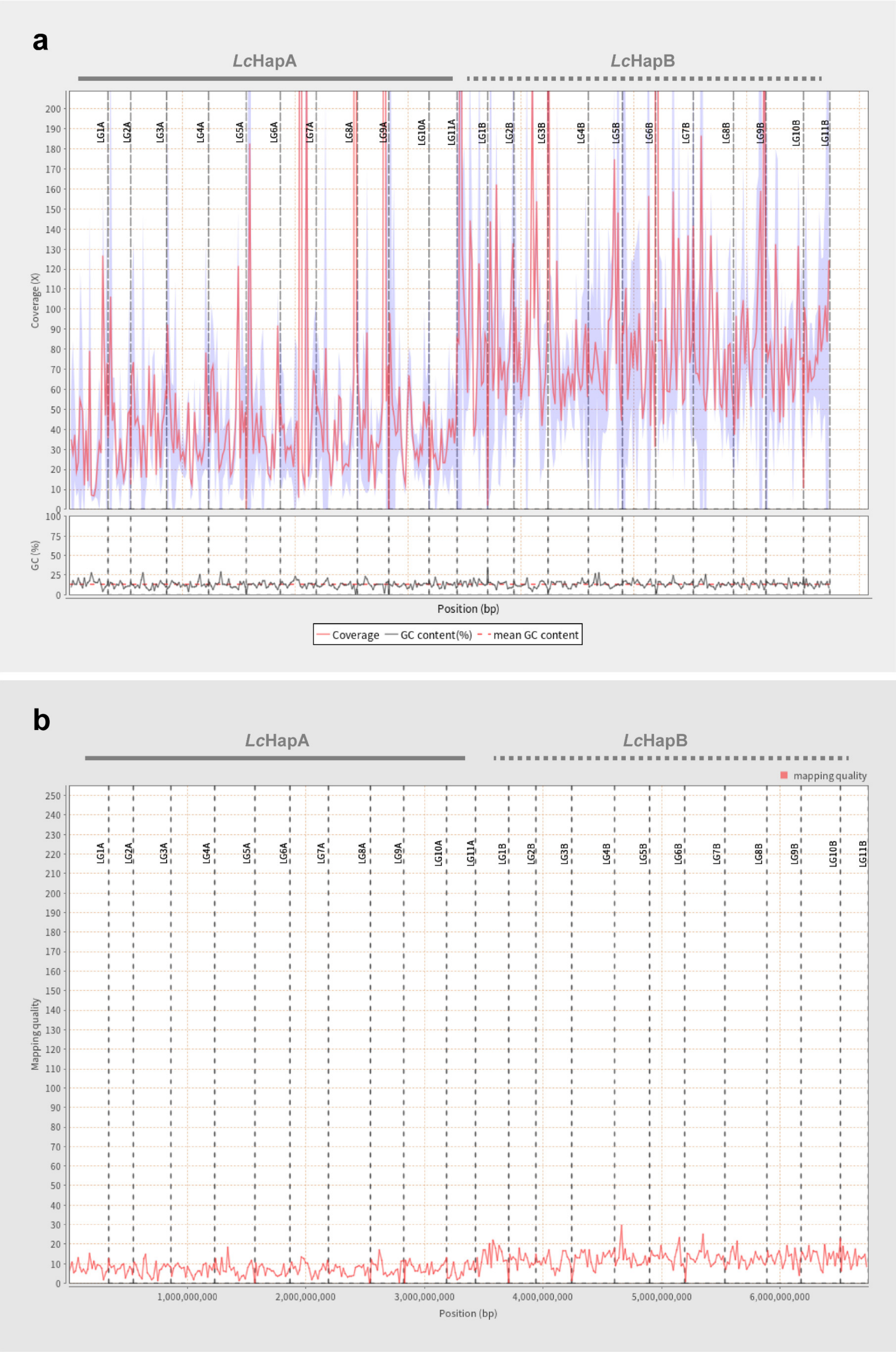


**Supplementary Figure 12. Mapping rate and mapping quality of the RNA-Seq reads of *L. sinense* onto genomes of *Lc*HapA and *Lc*HapB. a.** Mapping coverage and GC content. **b.** Mapping quality.

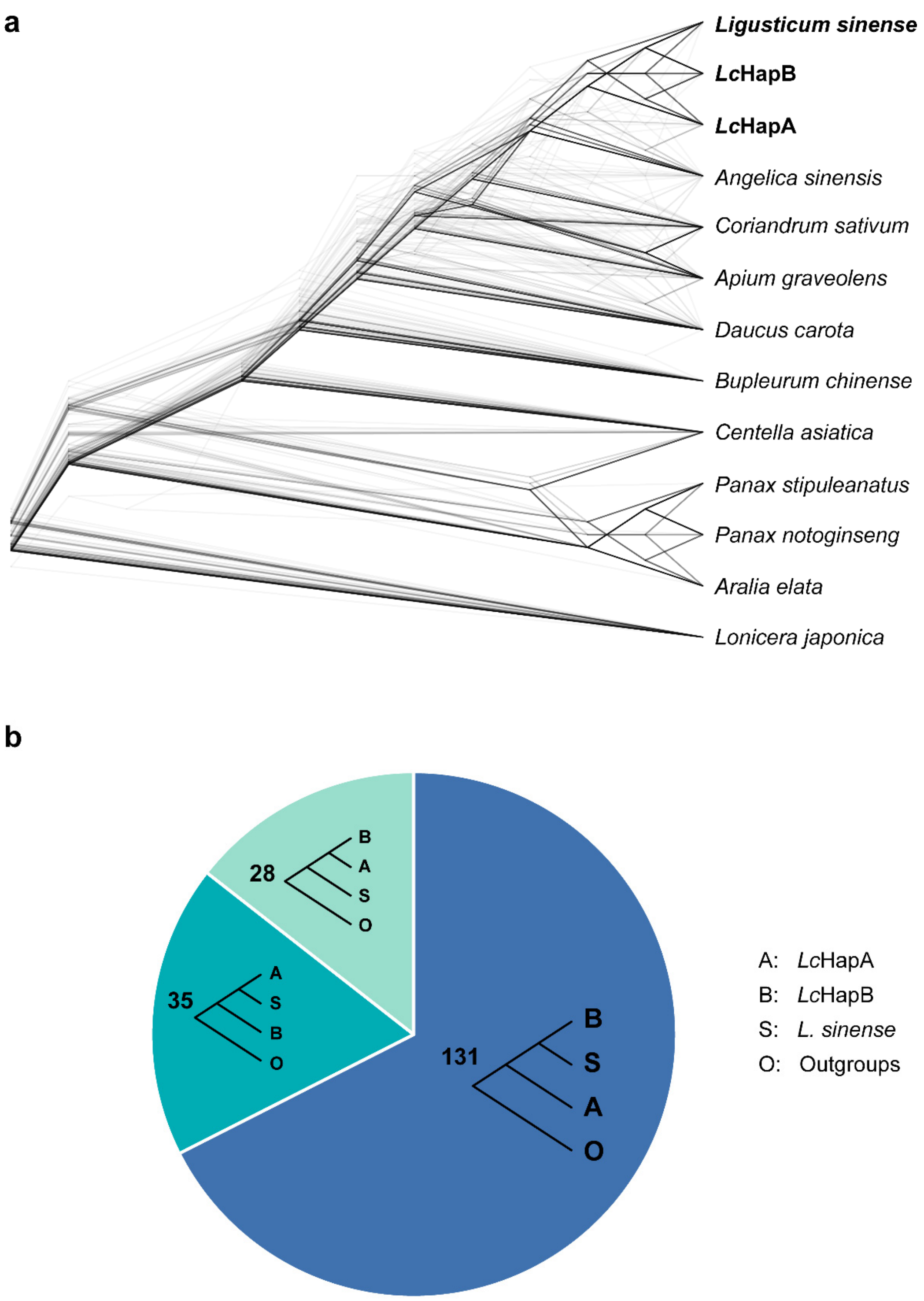


**Supplementary Figure 13. Summary of the possible phylogenetic topologies among *Lc*HapA, *Lc*HapB and other species. a.** Density gene trees base on 194 orthologous single-copy genes. **b.** Summary of the three possible topologies among *Lc*HapA, *Lc*HapB, and *L. sinense* based on the above 194 gene trees.

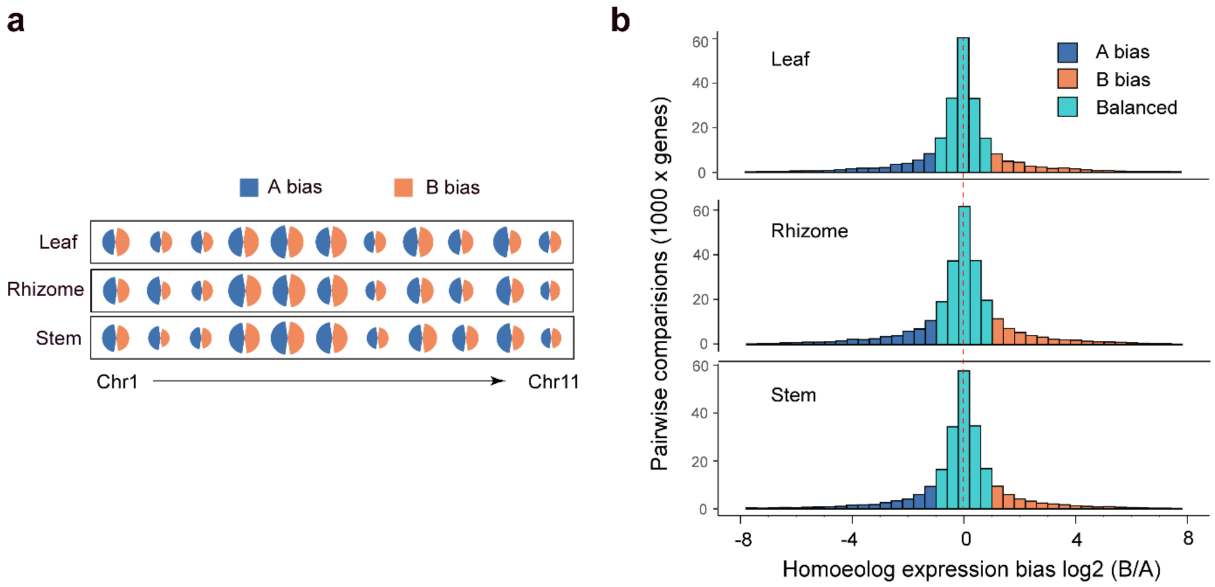


**Supplementary Figure 14. The distribution of ASEGs on different chromosomes and among tissues. a.** The relative percentage of ASEGs towards *Lc*HapA and *Lc*HapB in each chromosome. **b.** Distribution of ratio of expression abundance between homoeologous genes (*Lc*HapB / *Lc*HapA).

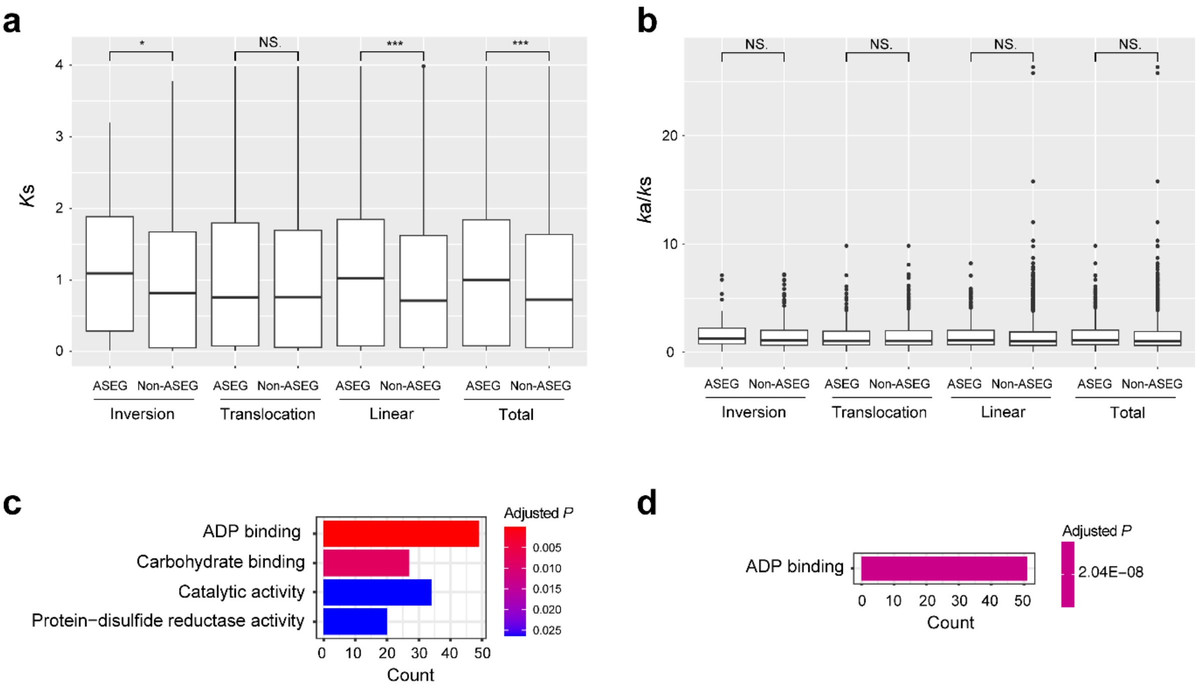


**Supplementary Figure 15. The comparisons for *Ks* and *Ka*/*Ks* ratio of ASEGs and non-ASEGs in *L. chuanxiong* and the GO enrichment of ASEGs. a.** Comparisons of *K*s between ASEGs and non-ASEGs in different SV types and across the whole genome. **b.** Comparisons of *K*a/*K*s ratio between ASEGs and non-ASEGs in different SV types and across the whole genome. **c.** GO enrichment for A-bias genes in at least one tissue. **d.** GO enrichment for B-bias genes in at least one tissue. * $P < 0.05$; ** $P < 0.01$; *** $P < 0.001$; NS., $P > 0.05$.

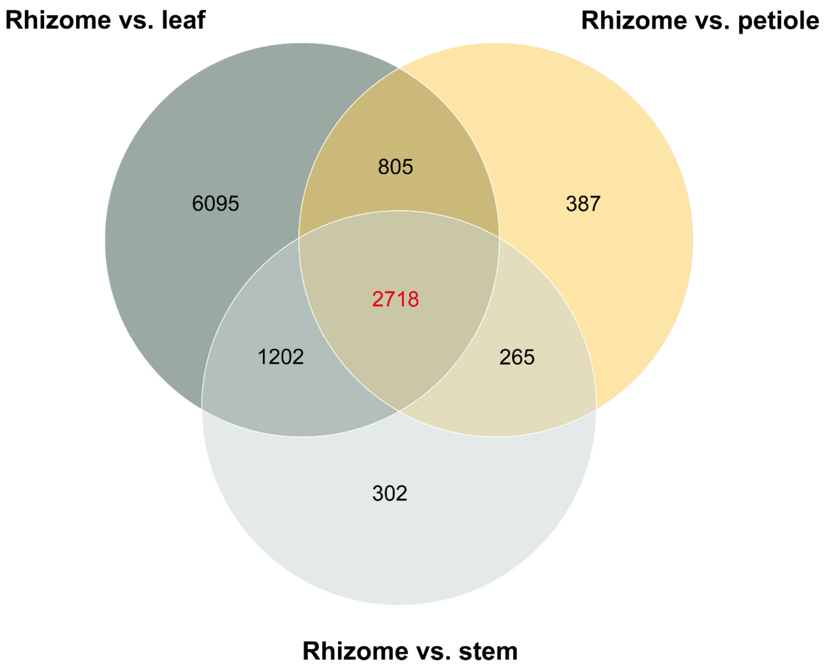


**Supplementary Figure 16. The Venn diagram of the number of significantly up-regulated genes with high expression (FPKM > 20) in rhizome compared to the other three tissues of *L. chuanxiong*.**

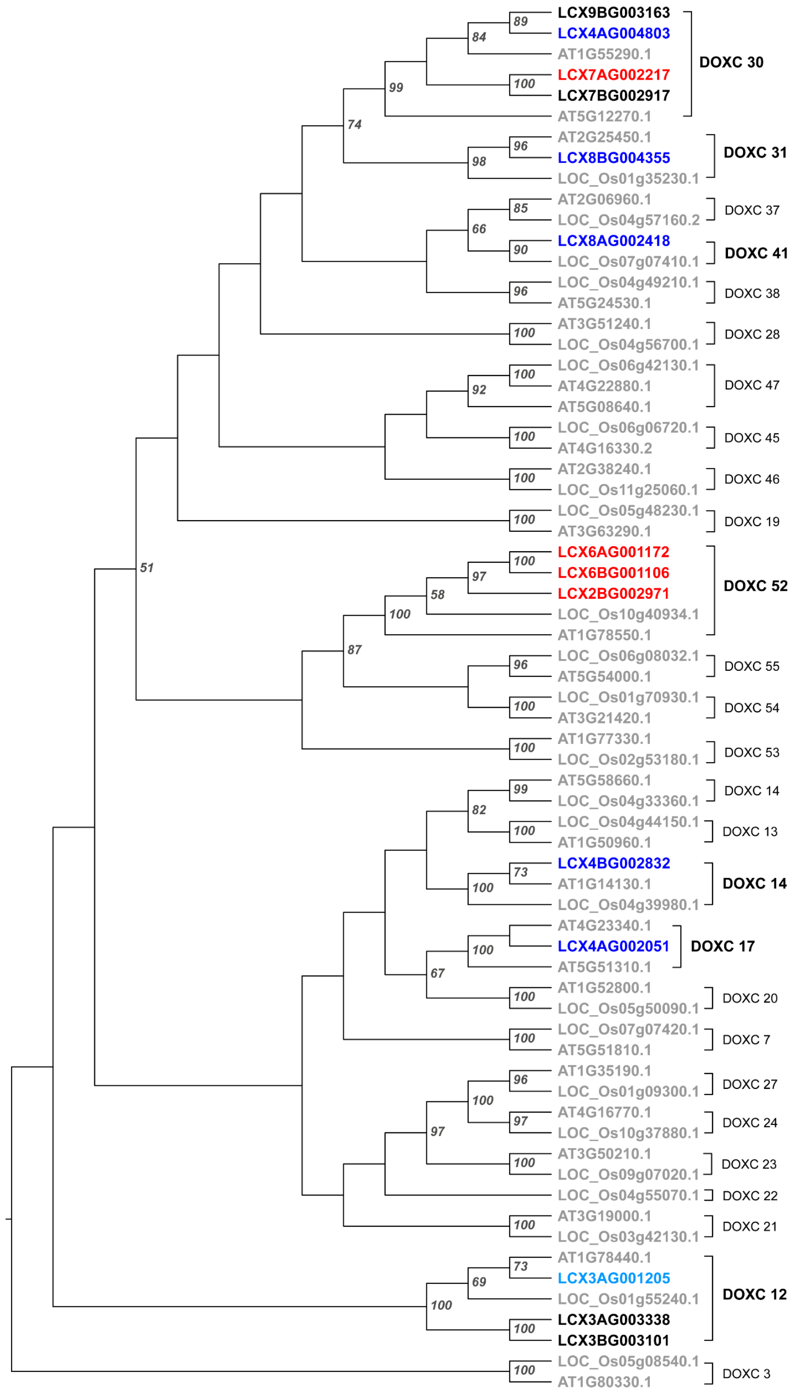


**Supplementary Figure 17. The ML phylogenetic tree of candidate 2OGD (DOXC family) genes for l-NBP (1) and butylidenephthalide (2) biosynthesis in *L. chuanxiong*.** The genes in grey represented 2OGDs (DOXC) in *Arabidopsis thaliana* and *Oryza sativa*, while genes in other colors represented 14 candidate 2OGDs in *L. chuanxiong* that belonged to seven DOXC clades. The genes in black were not cloned, those in light blue were failed to clone, those in blue were characterized but inactive for phthalide C-4/C-5 desaturation, and only those in red were active P4,5Ds.

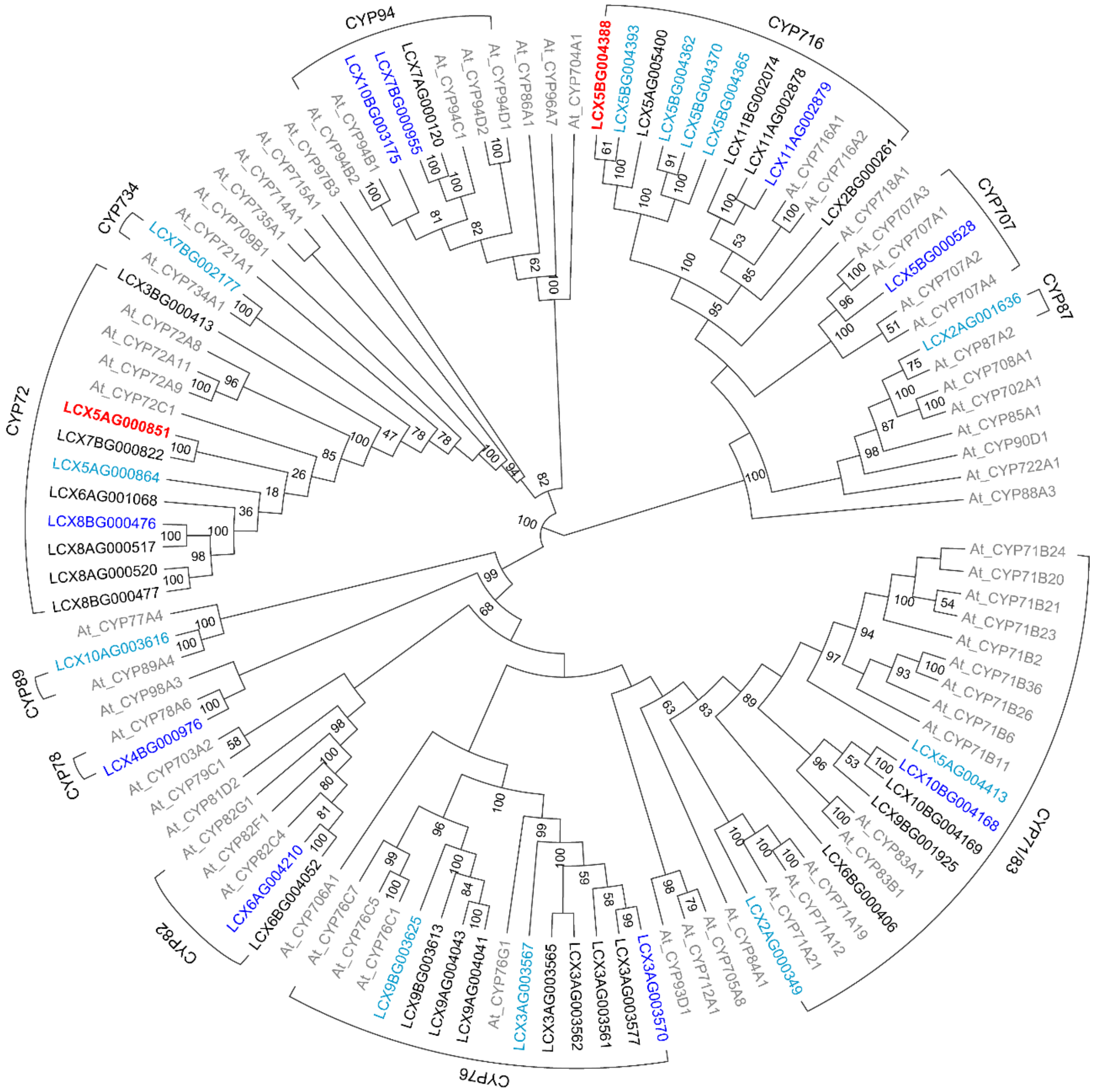


**Supplementary Figure 18. The ML phylogenetic tree of candidate CYPs for l-NBP (1) and butylidenephthalide (2) biosynthesis in *L. chuanxiong*.** The genes in grey represent CYPs in *A. thaliana*, while genes in other colors represent 45 candidate CYPs in *L. chuanxiong*, belonging to 12 CYP families. The genes in black were not cloned, those in light blue were failed to clone, those in blue were characterized but inactive for phthalide C-4/C-5 desaturation, and only those in red were active P4,5Ds.

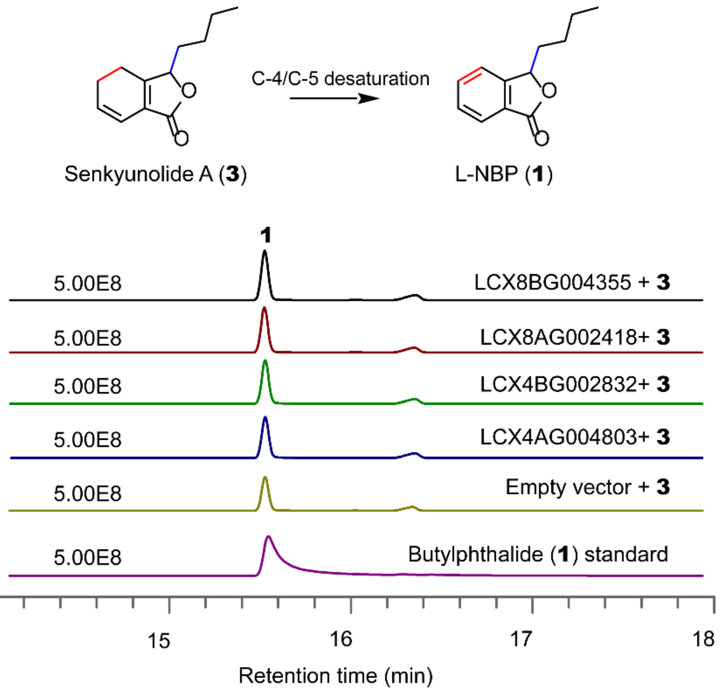


**Supplementary Figure 19. GC-MS peak plots of l-NBP (1) (*m/z* 133) in the indicated combinations of senkyunolide A (3) and candidate 2OGDs.** Except *Lc*2OGD1–4 (**Fig. 6**), described in the main text, the other candidate 2OGDs presented here failed to efficiently convert senkyunolide A (**3**) to l-NBP (**1**) in *E. coli*. Empty vector was used as negative control.

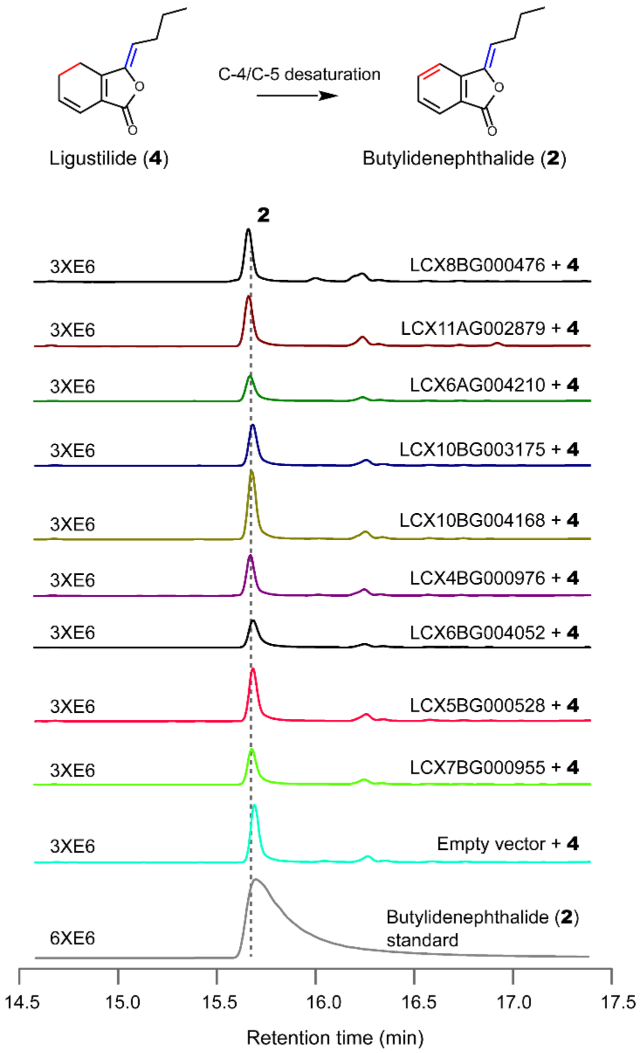


**Supplementary Figure 20. GC-MS peak plots of butylidenephthalide (2) (*m/z* 159) in the indicated combinations of ligustilide (4) and candidate CYPs.** Except *Lc*CYP716E94 and *Lc*CYP72A1132 (**Fig. 6**), described in the main text, the other candidate CYPs presented here failed to efficiently convert ligustilide (**4**) towards butylidenephthalide (**2**) in *N. benthamiana*. Empty vector was used as negative control.

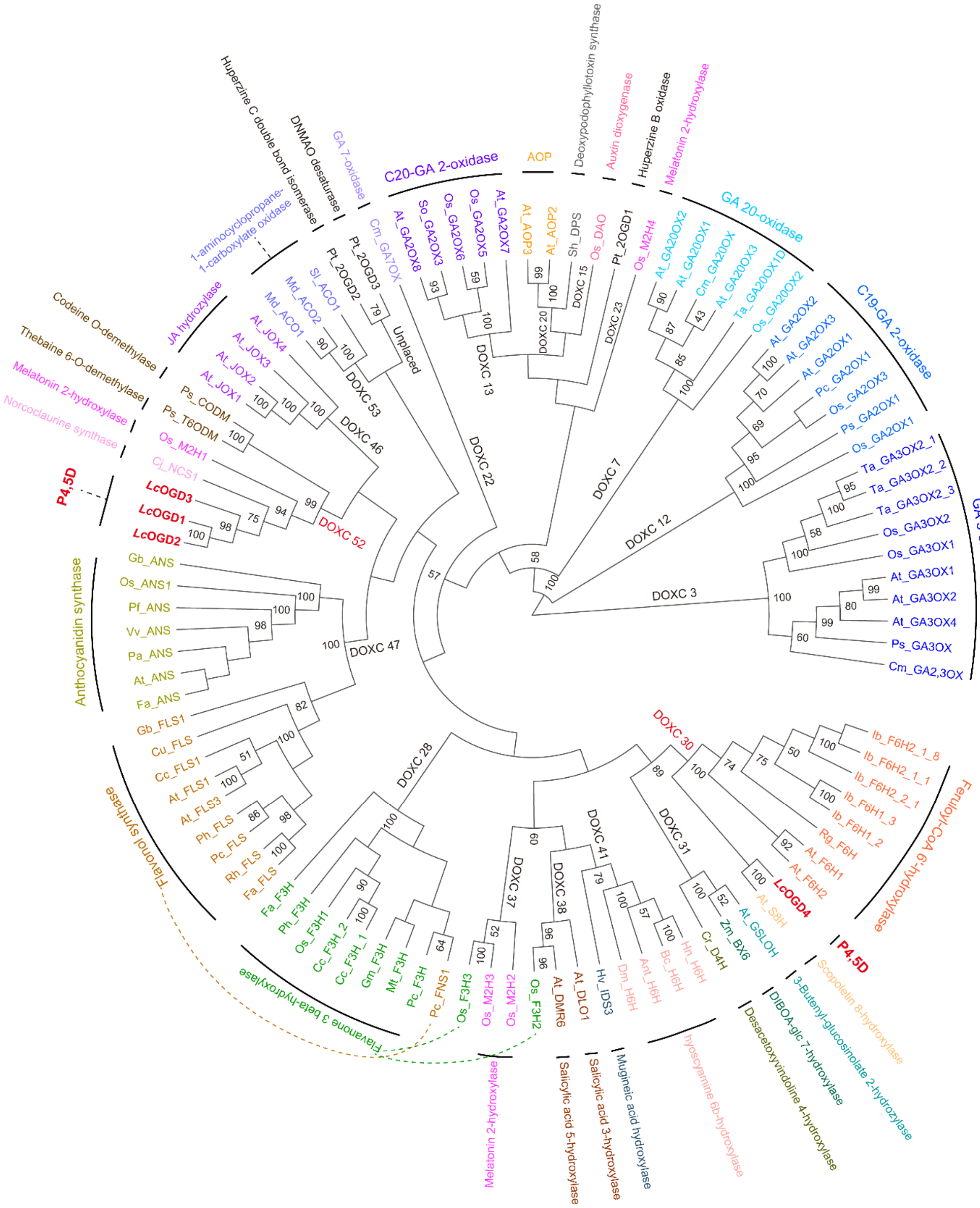


**Supplementary Figure 21. The ML phylogeny of characterized plant 2OGD (DOXC family).** The P4,5Ds are highlighted in red and bold.

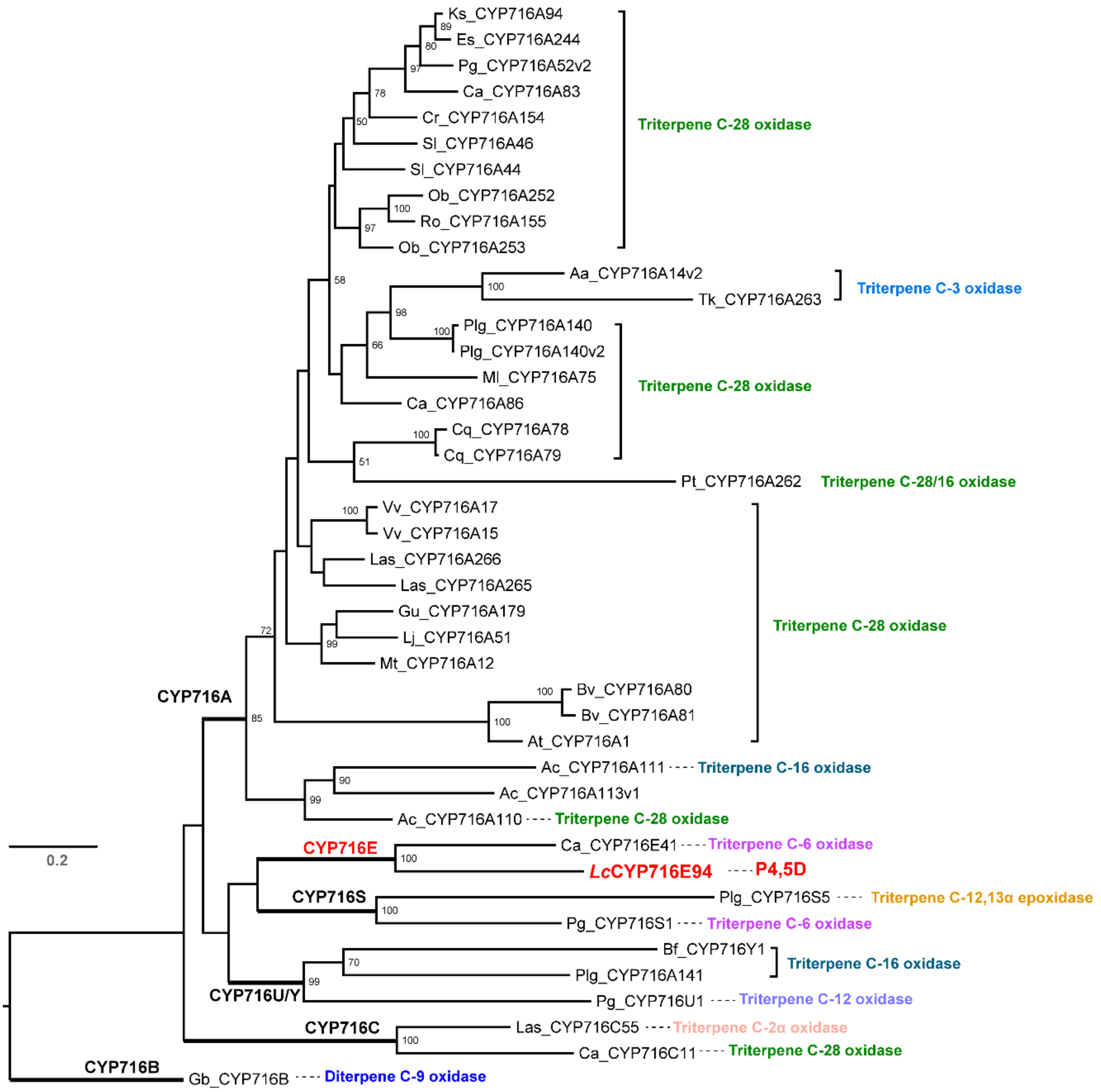


**Supplementary Figure 22. The ML phylogeny of characterized plant CYP716s.** The P4,5Ds is highlighted in red and bold.

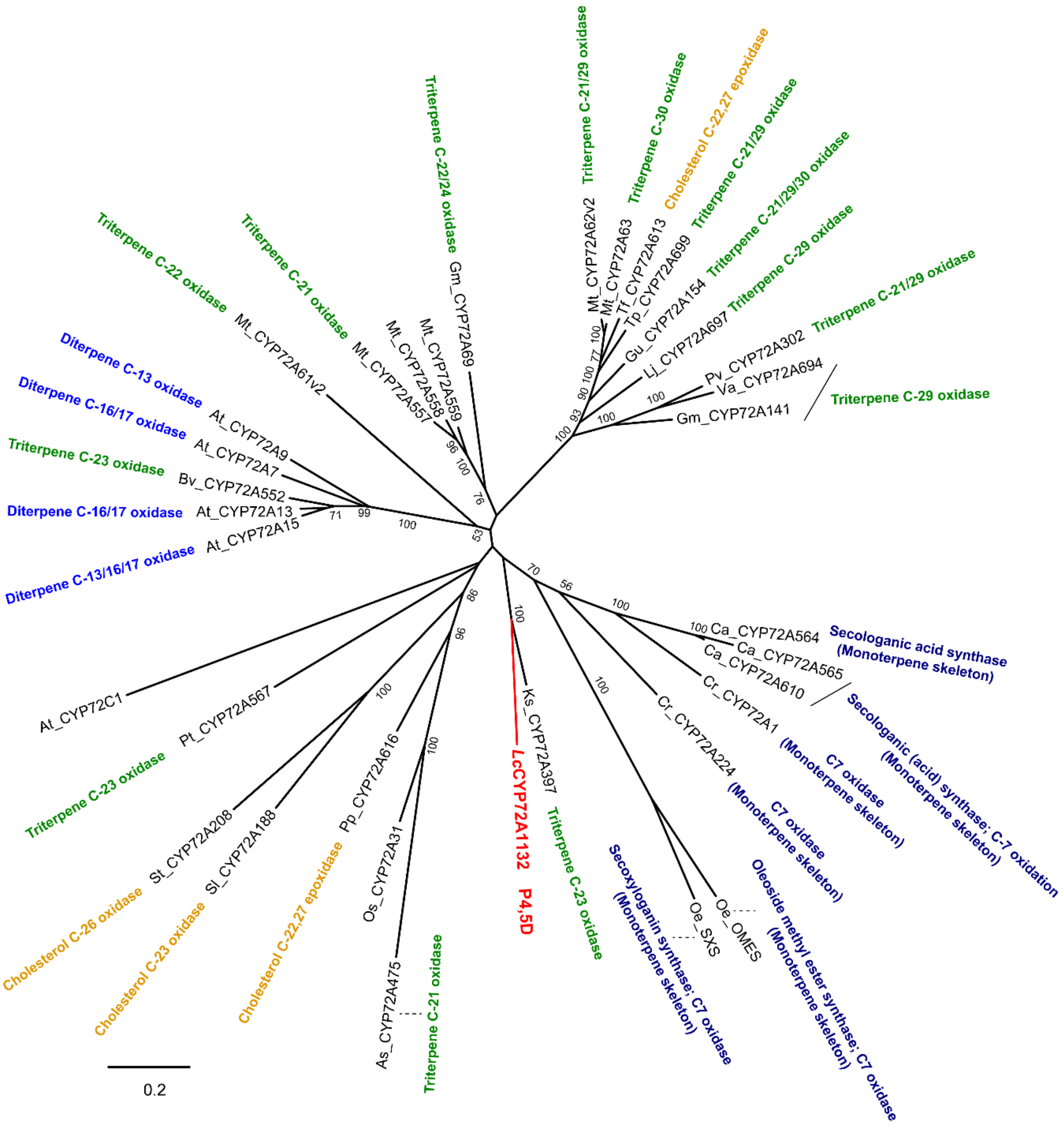


**Supplementary Figure 23. The ML phylogeny of characterized plant CYP72s.** The P4,5D is highlighted in red and bold.

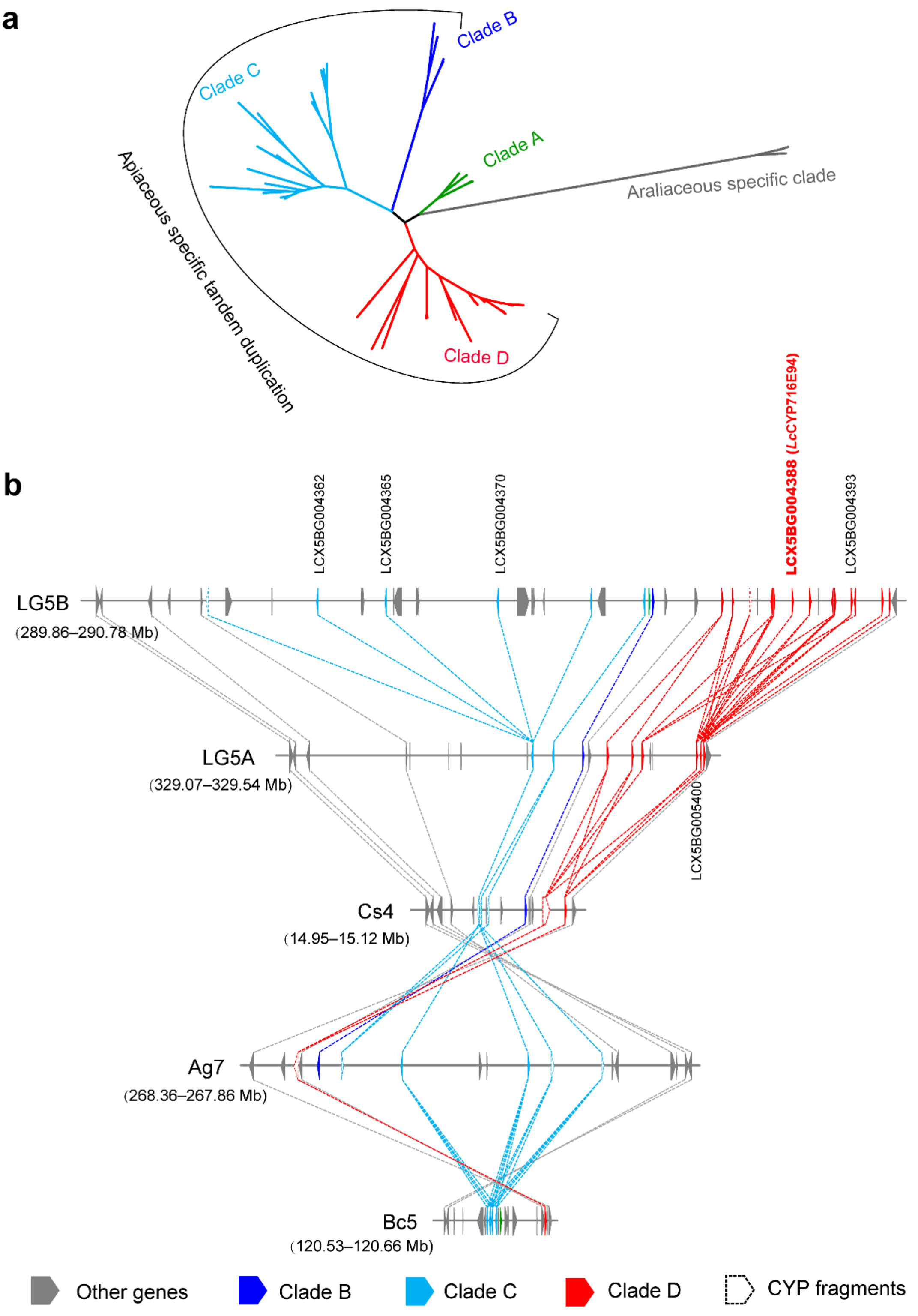


**Supplementary Figure 24. The tandem duplication of CYP716Es in Apiaceae. a.** The ML phylogeny of CYP716E subfamilies. The CYP716E subfamily in Apiaceae was subdivided into Clades A–D, among which Clades B–D were Apiaceae specific. **b.** The physical position of genes in Clades B–D in four Apiaceous species. LG5A/B, Chr 5 of *Lc*HapA/B; Cs4, Chr4 of *Coriandrum sativum*; Ag7, Chr 7 of *Apium graveolens*; Bc5, Chr 5 of *B. chinense*.

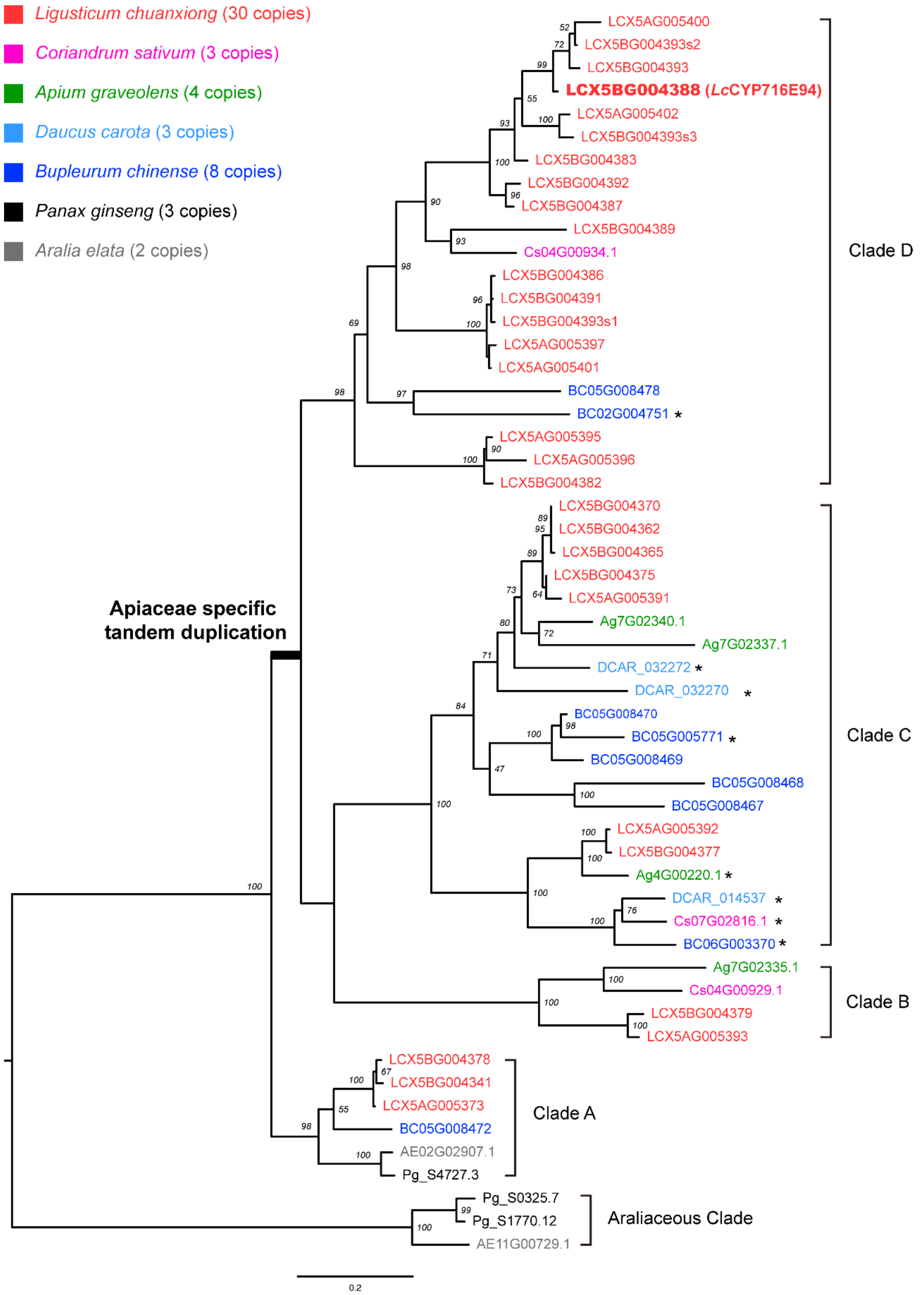


**Supplementary Figure 25. The ML phylogeny of Apiaceae specific tandem duplicated CYP716Es.** Genes marked with "*" are not located in the syntenic region of CYP716E tandem duplication.

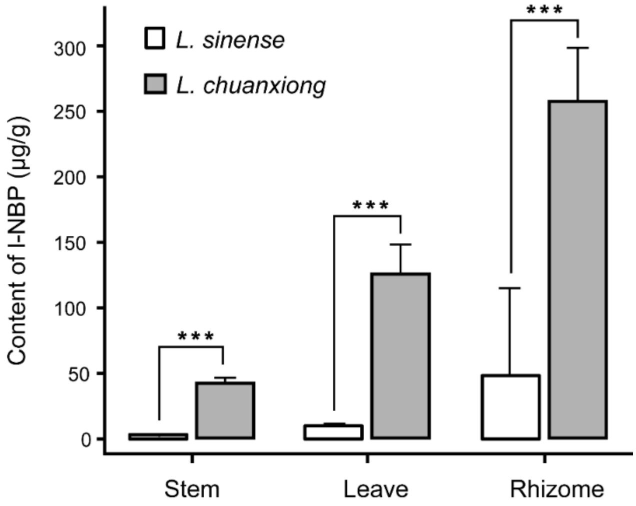


**Supplementary Figure 26. Comparison of the content of l-NBP between *L. chuanxiong* and *L. sinense*.** All content measurement were conducted for three samples and the error bars represent standard error of the mean. *** $P < 0.001$.

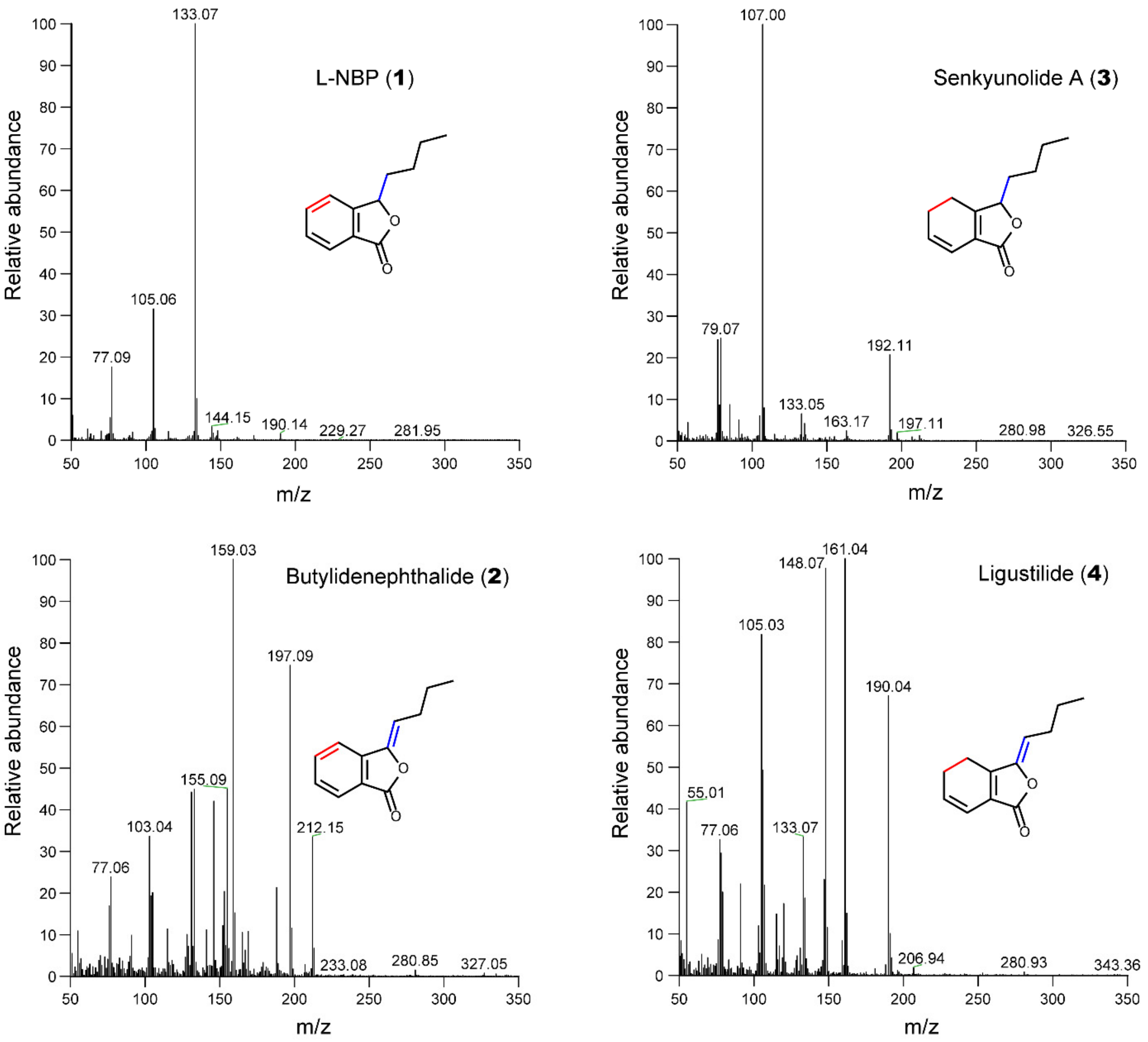


**Supplementary Figure 27. The mass spectrum of l-NBP (1), butylidenephthalide (2), senkyunolide A (3), and ligustilide (4).**
